# Generation and validation of homozygous fluorescent knock-in cells using CRISPR/Cas9 genome editing

**DOI:** 10.1101/188847

**Authors:** Birgit Koch, Bianca Nijmeijer, Moritz Kueblbeck, Yin Cai, Nike Walther, Jan Ellenberg

## Abstract

Gene tagging with fluorescent proteins is essential to investigate the dynamic properties of cellular proteins. CRISPR/Cas9 technology is a powerful tool for inserting fluorescent markers into all alleles of the gene of interest (GOI) and permits functionality and physiological expression of the fusion protein. It is essential to evaluate such genome-edited cell lines carefully in order to preclude off-target effects caused by either (i) incorrect insertion of the fluorescent protein, (ii) perturbation of the fusion protein by the fluorescent proteins or (iii) non-specific genomic DNA damage by CRISPR/Cas9. In this protocol^1^, we provide a step-by-step description of our systematic pipeline to generate and validate homozygous fluorescent knock-in cell lines.

We have used the paired Cas9D10A nickase approach to efficiently insert tags into specific genomic loci *via* homology-directed repair with minimal off-target effects. It is time- and cost-consuming to perform whole genome sequencing of each cell clone. Therefore, we have developed an efficient validation pipeline of the generated cell lines consisting of junction PCR, Southern Blot analysis, Sanger sequencing, microscopy, Western blot analysis and live cell imaging for cell cycle dynamics. This protocol takes between 6-9 weeks. Using this protocol, up to 70% of the targeted genes can be tagged homozygously with fluorescent proteins and result in physiological levels and phenotypically functional expression of the fusion proteins.

**Editorial Summary:** This protocol provides a detailed workflow describing how to insert fluorescent markers into all alleles of a gene of interest using CRISPR/Cas 9 technology and how to generate and validate homozygous fluorescent knock-in cell lines.

## INTRODUCTION

To study protein dynamics and functions within a cell, proteins are commonly tagged with fluorescent markers. Mahen *et al.*^1^ revealed that tagging proteins at the endogenous genomic locus is the best choice to study quantitative properties of the fusion protein by live cell imaging and biophysical methods. Cells generated and validated via this pipeline were used to study the nuclear pore assembly ^2^. Moreover, with the cell lines expressing homozygously tagged proteins using the described protocol it is possible to analyse the proteome dynamics in living cells ^3^ and systematically quantitatively map the cellular proteome in four dimensions using Fluorescence Correlation Spectroscopy -calibrated 4D live cell imaging ^1, 3^.

### Development of the protocol

Ran *et al.* ^5^ and Trevino *et al.* ^6^ have explained how to use Cas9 nickase to perform genome editing in eukaryotic systems. Our genome editing approach is based upon these genome editing methods using the paired Cas9 D10A nickase approach. We have used these methods to generate homozygously tagged genes of interest using a specific validation pipeline in a medium throughput manner as described in detail within this protocol.

All nucleases used for genome editing in mammalian cells (e.g. Zinc Finger Nucleases, Transcription Activator-like Effector Nuclease, CRISPR/Cas9) trigger a double strand DNA (dsDNA) break at a specific genomic locus which can be repaired by two major DNA repair pathways: non-homologous end joining (NHEJ) or homology directed repair (HDR) ^7^. To generate knock-in cell lines, HDR is essential to insert the tag of interest which is done in the presence of an exogenously introduced repair template during the dsDNA break ^5, 8–16^. CRISPR/Cas9 systems consist of either wildtype Cas9 or mutant Cas9 which perform either dsDNA breaks or nicking of single-stranded DNA, respectively, guided by small guide RNAs (gRNAs). The paired Cas9D10A nickase approach combines a Cas9 nickase mutant (D10A) with paired guide RNAs targeting the sense and antisense DNA strand of the target region to introduce targeted double-strand breaks ^5,6,10,17^ which are repaired by HDR in the presence of a DNA template containing the fluorescent marker. This approach is the most suitable method for generating endogenously tagged cell lines, as it avoids off-target effects while providing effective DNA repair for generating endogenously tagged cell lines ^6,16–20^ (Figure 1). We have routinely generated homozygous knock-in cell lines using the paired Cas9D10A nickase approach, which resulted in 70% of homozygous knock-in cell lines compared to 46% when using ZFNs. This data confirms that the paired Cas9D10A nickase approach is more susceptible to HDR as it was already shown by Bothmer *et al.*^18^ and Miyaoka *et al.*^19^, resulting in most likely in increased efficiency of homozygosity (please see “Comparisons to other methods” section for further details).

**Figure 1:**
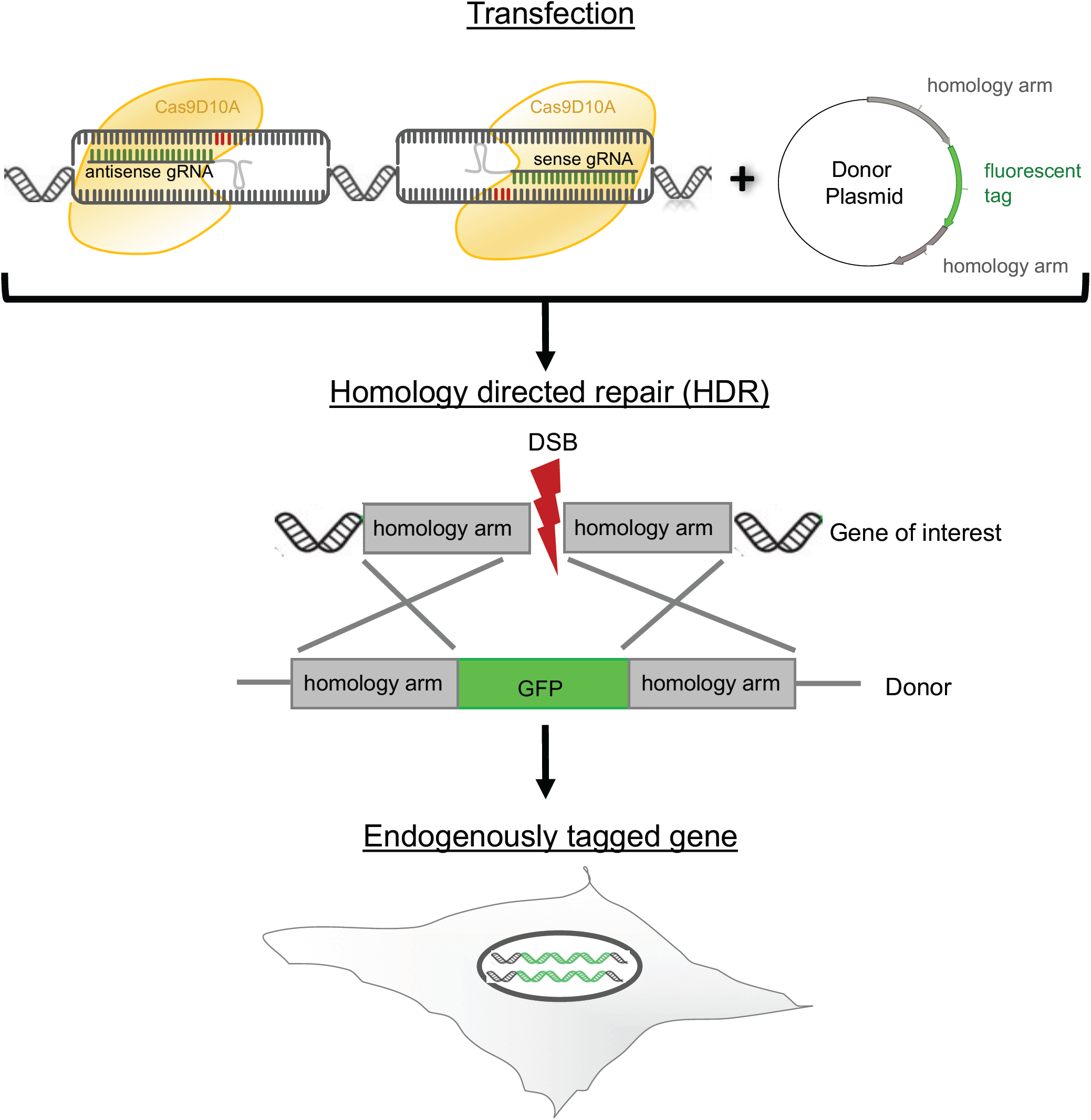
Paired CRISPR/Cas9 nickase approach. The RuvC nuclease domain is mutated in the Cas9D10A nickase resulting in single-stranded DNA breaks whose subsequent repair occurs preferentially through HDR ^6,14,17,18^. Two gRNAs are required to cleave the antisense and sense strand simultaneously when using Cas9D10A nickase which results in more specificity and less off-target cleavages^6,16–20^. Therefore Cas9D10A plasmids expressing nickase and either antisense or sense gRNAs are transfected in the presence of the DNA repair template, a donor plasmid containing the fluorescent marker flanked by homology arms (500-800 bp). This leads to a double strand break (DSB) followed by homology directed repair (HDR) and the fluorescent tag is inserted into a specific locus. The efficiency of HDR varies between cell lines and it is advisable to test the efficiency of homologous recombination in the cells of interest^19^.

There are several reports where inhibition of the NHEJ repair pathway has been used to increase HDR and subsequently homozygosity ^22–26^. However, we did not observe a significant improvement of homozygosity by depleting DNA Ligase 4 (LIG4) or X-ray repair cross-complementing protein 4 (XRCC4) via RNAi (unpublished data, B.K.). Also addition of Scr7, an inhibitor of the NHEJ repair pathway, did not increase the event of homozygosity which confirms the data of Greco *et al.*^26^ (unpublished data, B.K.). DNA damaging reagents can harm mitosis^27^ and because we mainly studied mitotic proteins, we did not want to disrupt the DNA repair pathway mechanism in case this affected the cell cycle.

There are several studies investigating DNA repair by NHEJ and HDR during the cell cycle in human cells ^28–30^. Homologous recombination is nearly absent in G1 phase, but most active in the S phase during the cell cycle ^30^. Consequently, we serum-deprived cells 24-48h prior to transfection, thereby synchronizing the cells partially in G1 phase. We then released the cells into the S phase during the targeted double strand DNA break to elevate the chance of HDR in the presence of the repair template. This approach lead to a minor increase of up to 2% of homozygosity for some gene targets but not all (unpublished data, B.K.). Although it is only a marginal increase of homozygosity, we performed serum-deprivation if possible because for some proteins of interest (POIs) only a few number of clones were homozygously tagged.

Homozygous tagging is not feasible for some essential proteins when structure and size of the FP tag perturb the function of the fusion protein. Therefore, some genes can only be heterozygously tagged. In our experience, and based on data in systematic cDNA tagging efforts, approximately 25% of human genes cannot be functionally tagged with FPs ^31^.

Careful validation of genome-edited cell lines is essential to control off-target effects which can be caused by insertion of the FP tag elsewhere in the genome, perturbation of the fusion protein by the FP, or non-specific DNA damage by unspecific binding of guide RNAs (gRNAs) leading to double strand DNA (dsDNA) breaks and genome rearrangements. Landry *et al.*.^32^ demonstrated that the genome of cell lines, such as HeLa Kyoto cells, is unstable which can lead to spontaneous mutations. Moreover, whole genome sequencing of every cell clone (up to 50 cell clones per candidate) is cost- and time-consuming. It is reasonable to consider whole genome sequencing with only a few validated cell clones when using non-cancer cell lines. The generation of many tagged cell lines is often necessary to study pathways or protein complexes and networks within a cell. Therefore, we have established a validation pipeline for genome-edited cell lines performing effective quality control at every level of the workflow (Figure 2) to ensure that high quality homo- and heterozygous cell clones of each POI are kept and used for further experiments. If only heterozygously tagged clones have been produced, a second round of genome editing of these clones is performed to achieve homozygosity. In addition, heterozygously tagged clones can have loss of function mutations in the untagged allele(s) due to dsDNA breaks without recombination, which can lead to pseudo-homozygous cell clones, expressing only the tagged fusion protein from the recombined allele(s). Details of each step within the workflow is discussed in the Experimental design.

**Figure 2:**
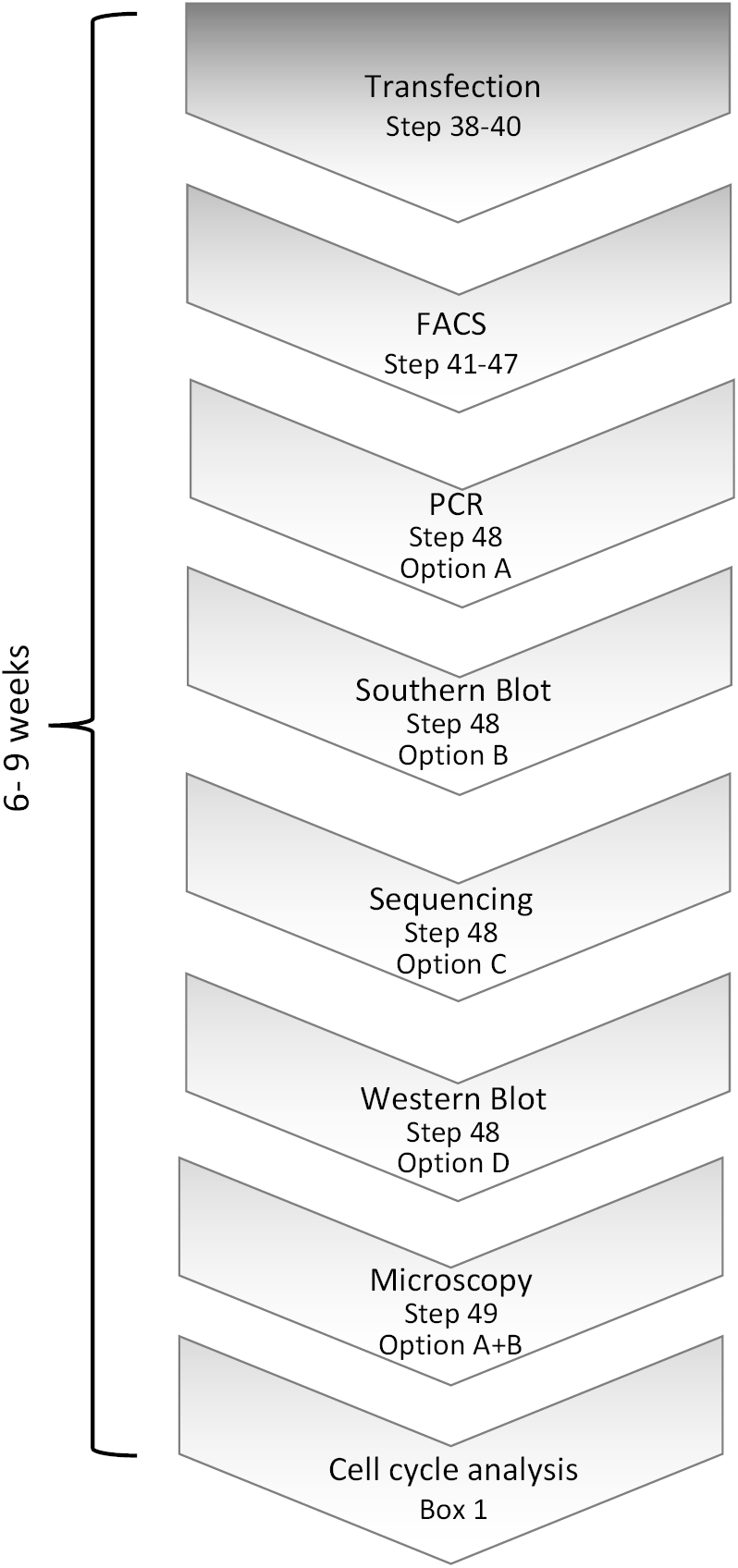
Validation pipeline of genome-edited cell lines. After the cells have been transfected and selected for fluorescently expressing clones, the validation pipeline consists of junction PCR, Southern blot analysis, Sanger sequencing, microscopy, Western blot and cell cycle analysis.

### Applications

It is possible to tag any POI using this protocol however, a major interest in our group is the characterization of proteins involved in cell division and in cell cycle control. Therefore, the protocol contains a cell cycle analysis step. If proteins of other signaling pathways are tagged, additional analysis might be necessary or the cell cycle analysis part can be exchanged. Endogenously tagged cells can be used for any applications whenever tagged proteins are needed that have to keep the physiological expression levels of the tagged protein and its functionality.

### Limitations

70% of the targeted genes were tagged homozygously with FPs and resulted in physiological levels and phenotypically functional expression of the fusion proteins using this protocol. Occasionally some proteins cannot be homozygously tagged because the structure and size of the FP tag perturb the function of the fusion protein. Therefore some genes can only be tagged heterozygously. Moreover, how many molecules of the POI are expressed should also be considered because there are detections limits for some applications when expressing low amounts of tagged proteins.

### Comparisons to other methods

CRISPR/Cas9 technology is developing rapidly. One of the major issues are off-target effects, which can be significantly decreased using the paired Cas9D10A nickase approach. In addition, transfection of pre-formed Cas9 gRNA complexes (Cas9 ribonucleoprotein = Cas9RNP) has been reported to increase the efficiency of gene knock-out with less off-target effects ^33–35^. However, in our experience using Cas9RNP or mRNA rather than plasmids to express Cas9 protein does not improve the efficiency of homozygous knock-ins in human cell lines (unpublished data, BK) but rather results largely in heterozygously tagged genes. Roberts *et al.*^36^ also reports only of heterozygously tagged genes in their systematic gene tagging approach using CRISPR/Cas9 RNP in human stem cells. However, given that the application of this protocol is to then perform quantitative FCS-calibrated live cell imaging (see our accompanying Nature Protocol Politi *et al.*)^4^. This is dependent on homozygous fluorescent tagged proteins, therefore the Cas9RNP approach with its current performance is not suitable for this pipeline at the moment.

Paix *et al.* ^37^ has demonstrated that linear DNAs with short homology arms used as a DNA repair template can be applied for genome-editing knock-in approaches. We were able to use double stranded linear DNA templates consisting of mEGFP flanked by short 50 bp homology arms, but could only generate heterozygous knock-in cell clines, without the production of homozygous knock-ins (unpublished data, B.K.).

Paquet *et al*. ^38^ has shown that the homozygous introduction of mutation using single-stranded oligo donor (ssODN) via Cas9RNP requires a gRNA targeting close to the intended mutation. In our protocol the gRNAs are also placed as close or even over the target site at which the tag should be introduced consequently resulting in a good efficiency of homozygous introduction of fluorescent markers or other tags, such as Halo or SNAP, at the locus of interest. However, we need to introduce a fluorescent tag via plasmid donors that is around 700-750 bp and consequently we cannot use ssODNs, the approach described by Paquet *et al.*2016^38^. As discussed above the Cas9RNP approach in combination with plasmid DNA as donor was in our hands not successful for homozygous tagging, although heterozygous tagging was possible. In summary, the paired Cas9D10 nickase plasmid based approach described in this protocol remains the optimal procedure for the generation of human cell lines with homozygous fluorescent gene knock-ins at the moment.

## Experimental design

### Design and delivery of gRNAs and donor plasmids

The design of gRNAs and donors depend on where the POI should be tagged, typically either at the N-or C-terminus. It is prudent to perform an extended literature search for localization data, functionality studies of tagged versions of the POI and expression data in other species such as yeast or mouse to provide insight as to which end of the POI can be tagged while maintaining its functionality intact and localization correct.

Point mutations within the seed region, which is PAM-proximal 10-12 bases, can have decreased gRNA binding activity^39^. Therefore sequence variations such as single nucleotide polymorphisms (SNPs) of the used cell line, which differ from the reference genome, and isoforms of the GOI can interfere with CRISPR/Cas9 targeting. Consequently, it is advisable to perform sequencing not only within the region of the GOI used for tagging but also of possible isoforms.

The gRNAs should bind directly to the target site, such as the start or stop codon of the open reading frame. The probability of HDR is best within 100 bp of the double strand DNA break site. Beyond that the efficiency of HDR drops by four fold ^9^. Two gRNAs, one for the antisense and one for the sense DNA strand, are necessary when using the Cas9D10A nickase. Appropriately designed gRNAs ^6,9,17,18^ are inserted as annealed oligonucleotides into Cas9/gRNA expression plasmids containing two expression cassettes, a human codon-optimized Cas9D10 nickase and the guide RNA.

The donor plasmid contains the template for homologous recombination with the fluorescent marker gene (e.g. mEGFP or mCherry) flanked by 500-800 bp homology arms complementary to the target site of the GOI. The fluorescent marker replaces either the start or the stop codon and is in-frame with the GOI. By default, there should be a short flexible linker incorporated between the tag and the gene to improve functionality of the tagged protein. Neither an exogenous promoter nor a mammalian selection marker should be part of the donor plasmid to prevent expression of the marker gene without recombination with the target gene.

The delivery method of the plasmids depends on the cell line of choice and should be optimized. HeLa Kyoto cells are transfected very well using lipofection which is used for the generation of endogenously tagged cell lines within this protocol.

### Fluorescence activated cell sorting of Fluorescent Protein positive cell clones

Donor plasmids are used as DNA repair templates. BacORI is essential to replicate plasmids within *E.coli*, but contains cryptic promoter regions that can cause some background expression in transfected cells prior to genomic integration for as long as the plasmid is maintained ^40^. Therefore, single cell sorting to select fluorescent cells is performed 7-10 days post transfection reducing the number of false positive cell clones during Fluorescence activated cell sorting (FACS). Depending on the survival rate of single cells of the cell line of choice after FACS, several hundreds of single cell clones should be sorted for downstream characterization. Often, the best growth condition for single cell clones has to be optimized for each parental cell line. The conditions in this protocol have been optimized for widely used HeLa Kyoto cells. HeLa Kyoto wt cells were used as negative control to distinguish between non-specific and specific FP/mEGFP expression levels.

Cells were sorted with the DAKO MoFlo High Speed sorter and performed by the Flow Cytometry Core Facility at EMBL (Heidelberg, Figure 3). The settings and calibrations are as follows: BD FACSFlow ™ was used as sheath fluid. The instrument was fitted with a 100-μm nozzle tip and the frequency of droplet generation is around 40 kHz. The primary laser was a Coherent Argon ion Sabre laser, tuned to single line 488 nm 200 mW output power (100 mW hitting the cells after passing through a 50/50 beam splitter). This laser was used in the generation of Forward and Side scatter as well as in the excitation of GFP. The secondary laser was Coherent Sabre Krypton ion laser. It was tuned either to single line 568 nm (200 mW) in the excitation of mCherry or to Multiline Violet (MLVio: 406.7 - 415.4 nm), in the excitation of Cerulean. Laser beams were carefully aligned to the MoFlo’s primary optical path. The alignment of the optical path is optimized while running Flow-Check ™ Fluorospheres. Sample events are acquired at less than 25% of droplet occupancy. The sample-sheath differential pressure was low to confine cells at the centre of the co-axial flow, thus limiting illumination variation. Sort decision gates were based on a combination of regions drawn around target populations in several 2-D plots: FSC-Area vs. SSC-Area; FSC-Area vs. FSC-Pulse Width and fluorescence. A Single 1-drop mode was used during the cloning of sort classified target cells into 96-well plates containing growth media. Data was analyzed using Moflo Summit software and FlowJo (Tree Star, Inc.) software.

**Figure 3:**
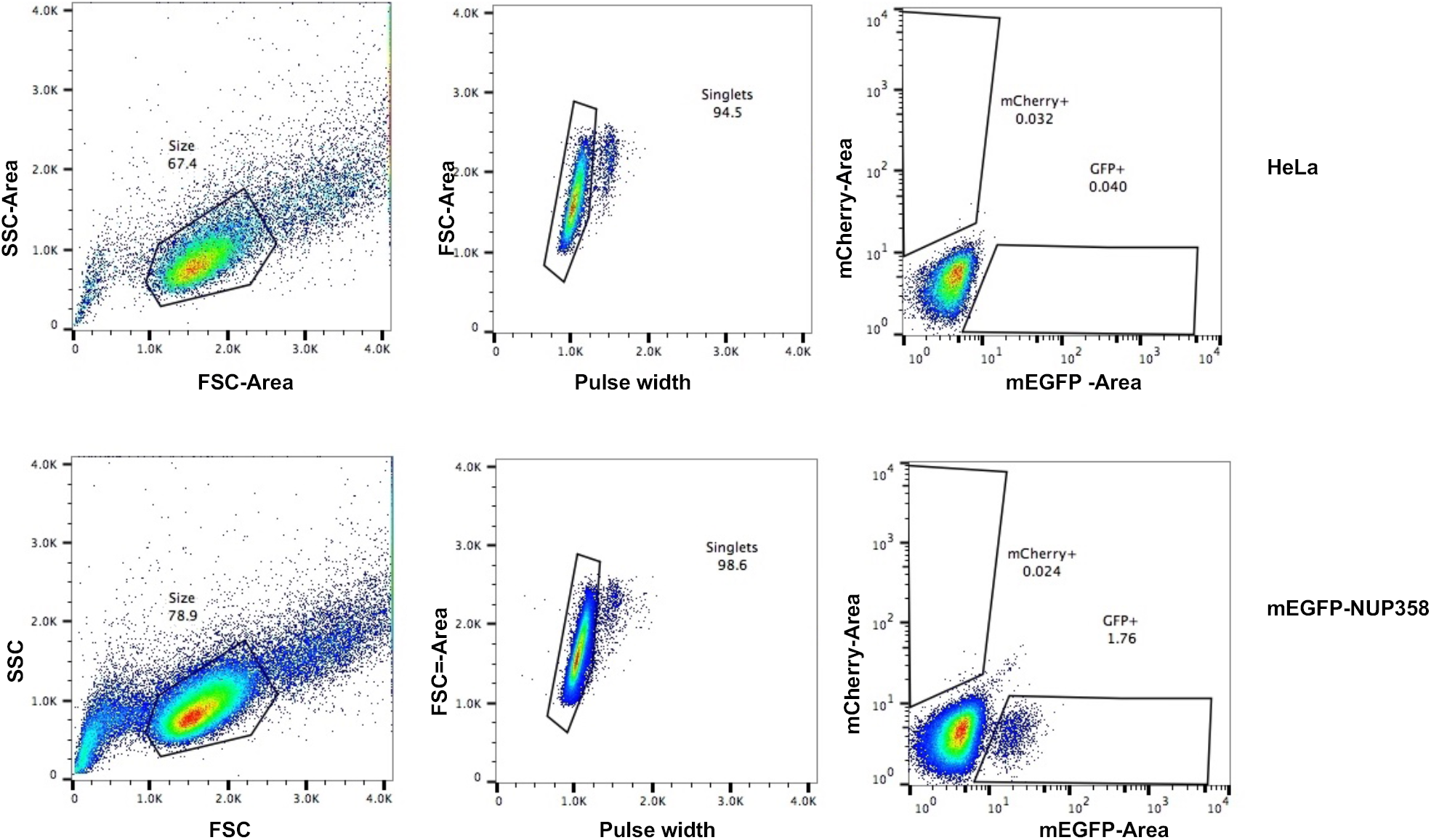
FACS of genome-edited cell lines tagging NUP358 at the N-terminus. HeLa Kyoto cells are used as negative control and mEGFP-NUP358 cells were selected for mEGFP expression. GFP-expressing single cells were sorted into 96well plates using a DAKO MoFlo High Speed sorter. The panels on the left display the side scatter area vs. forward scatter area to gate for live cells (black polygon, number = % of live cells). The panel in the middle distinguishes singlets (black polygon, number = % single cells) using the FSC-Area vs. FSC-Pulse Width plot. The diagrams on the right exhibit the signal intensity of mEGFP using a 512/15 bandpass filter and the red polygon indicates selected cells for single cell sorting (number = % of GFP positive cells). HeLa Kyoto wt cells serve as a negative control and define background signal and the gates for the positive GFP cells were set accordingly. 1.76% mEGFP-NUP358 positive cells were sorted into 96-well plates to obtain single clones expressing mEGFP-NUP358. SSC = Sideward Scatter, FSC = Forward Scatter.

The fluorescent signal of endogenously tagged proteins depends on its physiological expression level and is often relatively dim compared to commonly used overexpression systems and high sensitivity FACS sorters need to be used. For cell cycle regulated proteins, it might additionally be necessary to synchronize cells in a specific cell cycle stage to enrich cells with high signal for FACS. The efficiency of positively tagged cells 7-10 days after plasmid transfection ranges between 0.05-10% (e.g. Figure 3), but is typically around 0.5%. HeLa Kyoto wt cells serve as a negative control and define background signal (Figure 3, upper panel Hela cells, 0.04% GFP-positive) and the gates for the positive GFP cells were set accordingly. In the case of mEGFP-NUP358, 1.76% mEGFP-NUP358 positive cells were obtained (Figure 3, lower panel, mEGFP-NUP358) which were sorted into 96-well plates.

### Junction PCR

Correct integration of the fluorescent marker into the expected locus can be tested by PCR, using primers binding outside of the homology arms in the genomic locus and within the recombined fluorescent tag (Figure 4A). To determine if all alleles are tagged with the fluorescent marker (= homozygosity), primers located outside of the right and left homology arm are used. The resulting one or two PCR products indicate homozygosity or heterozygosity, respectively. This is not a quantitative PCR, therefore the tagged-gene in heterozygous clones might not be detected due to competition between the two PCR products (Figure 4B). Consequently, it is advisable to repeat the junction PCR with primers within the homology arms resulting in shorter PCR products (Figure 4C) improving amplification of both PCR products. However, homozygously tagged genes will always result in only one PCR product when using primers for the GOI at the target site (Figure 4, clone #97).

**Figure 4:**
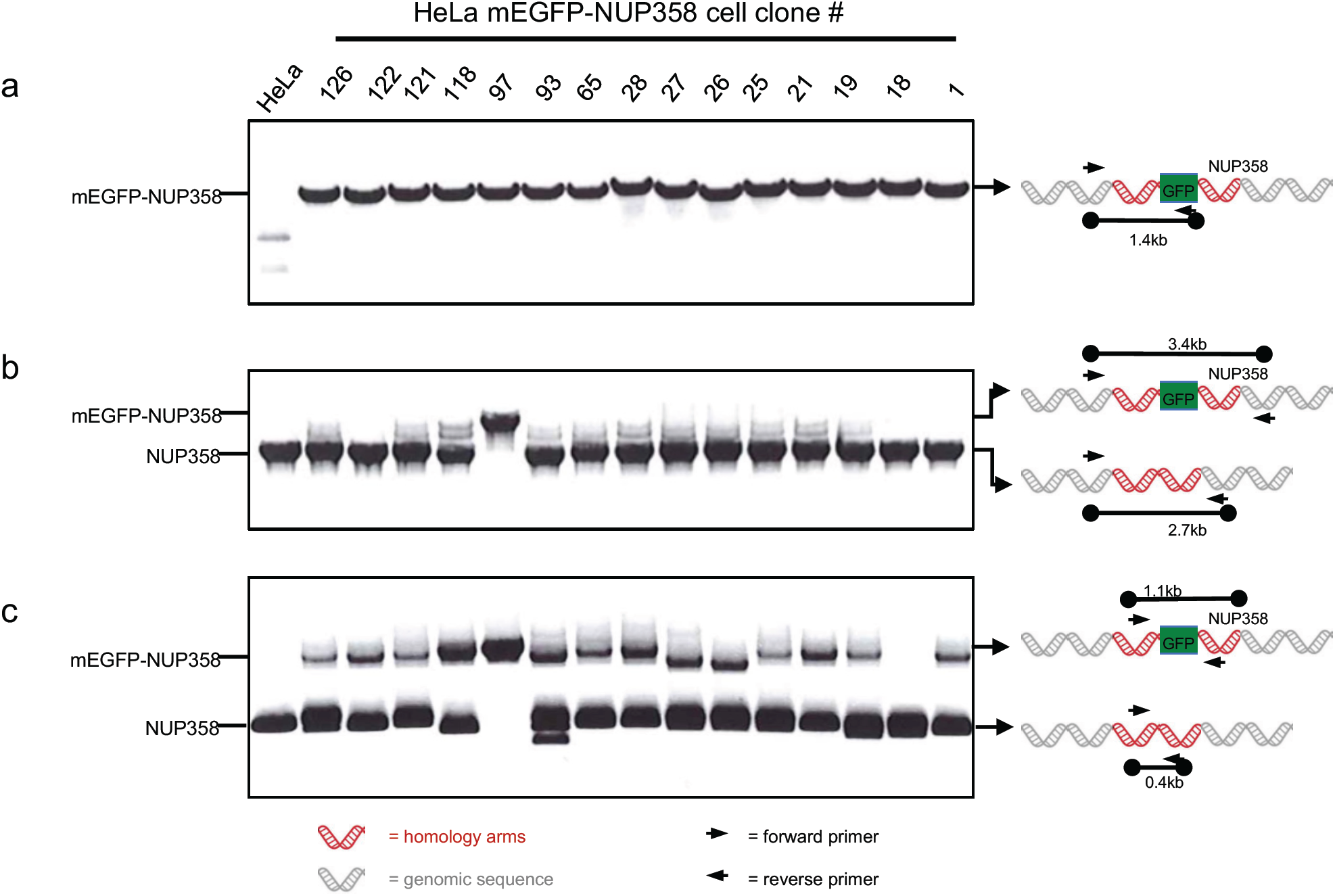
Junction PCR of HeLa Kyoto mEGFP-NUP358 cells. The junction PCR was performed as described in steps 48 Option A. (a) PCR to test the integration of mEGFP into the correct locus. The forward primer binds at the 5’ end outside of the left homology arm and the reverse primer to the fluorescent marker gene (see Table 1) resulting in a single 1.4 kb fragment if NUP358 is tagged at the N-terminus with mEGFP. The binding of the primers is illustrated in the scheme at the right hand side. (b) To test if all alleles are tagged with the fluorescent marker at the correct locus (homozygosly tagged mEGFP-NUP358), a forward primer located 5’ outside of the left homology arm and a reverse primer 3’outside of the right homology arm were used. As depicted in the scheme two bands can occur: either untagged Nup358 as 2.7 kb fragment or mEGFP-tagged NUP358 as 3.4 kb fragment. Homozygosity is indicated when only the tagged mEGFP-NUP358 (3.4 kb fragment) is detected as demonstrated for clone #97. (c) Additional PCR to test for homozygosity using primers within both homology arms of the donor (see design of primers for T7E1 assay) because problems occurred in detecting the tagged gene as demonstrated in (b). The forward primer is within the left homology arm whereas the reverse primer is located in the right homology arm. Two fragments can be detected: untagged Nup358 results in a 0.4 kb fragment and tagged mEGFP-NUP358 in a 1.1 kb fragment. Only the tagged mEGFP-NUP358 fragment of 1.1 kb can be detected for the homozygous clone #97.

### Sanger sequencing

PCR performed with primers around the target region followed by Sanger sequencing ^41,42^ is used to detect mutations within the insertion site of the tag at the genomic locus. Sequences of parental wildtype cells and the donor plasmid are used as reference. A sequencing example followed by alignment analysis is shown in Figure 5. Two different HeLa Kyoto cell clones either expressing mEGFP-NUP358 homozygously (Figure 5, #97 tagged) or heterozygously (Figure 5, #118 tagged and #118 untagged) are aligned to the mEGFP-NUP358 donor plasmid and HeLa Kyoto wt sequences. HeLa Kyoto wt and #118 untagged do not contain mEGFP as expected, but there is a deletion (63nt) within the #118 untagged NUP358 gene causing a deletion of 6 amino acids in the first exon of the untagged NUP358 (Figure 5, #118 untagged). However, the mEGFP-NUP358 of the same clone (Figure 5, #118 tagged) is correct and mEGFP is integrated without any mutations at the insertion sites (Figure 5, #118 tagged) as expected. The homozygote clone #97 contains only the tagged mEGFP-NUP358 whereby the mEGFP is inserted correctly into the target site (Figure 5, #97 tagged). Consequently, the mEGFP was correctly inserted at the N-terminus of NUP358 and this homozygously tagged clone can be used for further experiments.

**Table 1:**
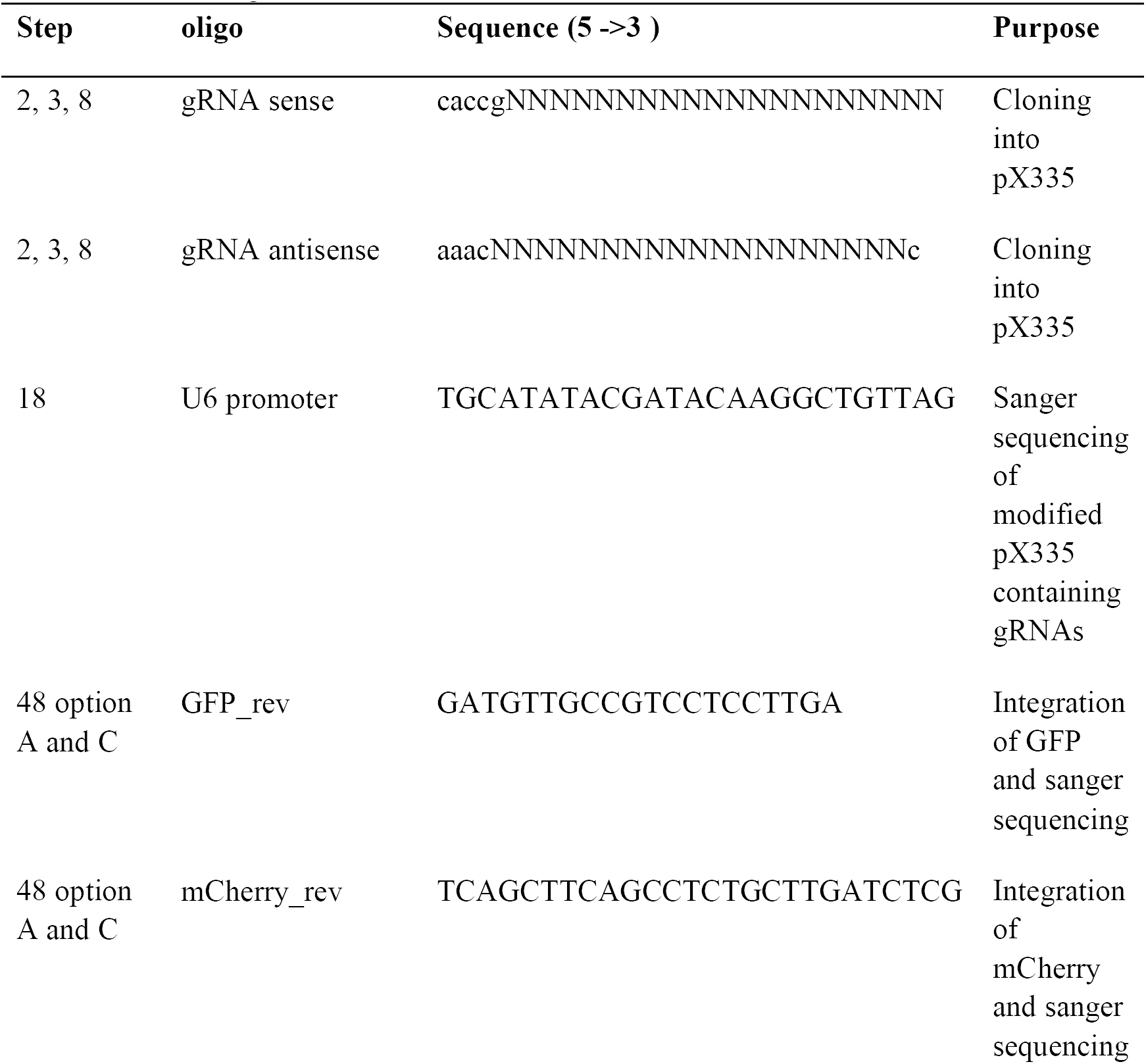
Oligos

**Figure 5:**
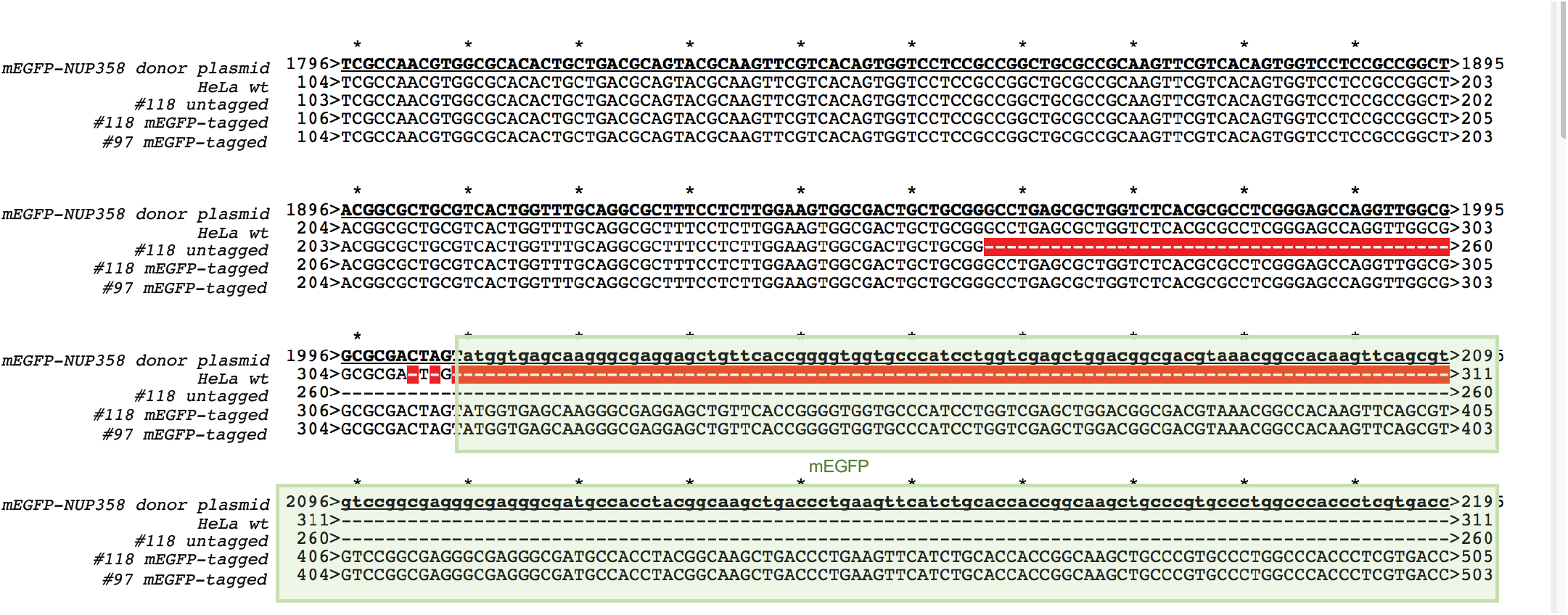
Sanger sequencing at the target site (Start codon). PCR was performed using primers around the target site, start codon of NUP358, followed by sanger sequencing. The sequences of tagged and untagged NUP358 was aligned to the donor plasmid used as DNA repair template during transfection. The box indicates mEGFP which is missing in HeLa Kyoto wt and untagged NUP358 as expected. The red line within the diagram depicts missing nucleotides (= deletion).

Additionally, if there are only heterozygous cell clones after the first round of genome editing, all untagged alleles of heterozygous cell clones are sequenced to detect possible mutations within the target region. This is essential since point mutations could hinder gRNA binding in the second round of genome-editing required to generate homozygous cell clones. Some proteins cannot be homozygously tagged due to the structure and size of the FPs which can disturb the function of the protein.

### Southern blot analysis

Southern blot analysis is important to show homozygosity and to exclude off-target integration of the tag. A probe against the tag should result in only one expected product on the Southern Blot (Figure 6, GFP probe, mEGFP-NUP358). However in some cases additional DNA fragments are detected indicating either random integration of the donor plasmid or other off-target effects due to genome rearrangement after double strand DNA breaks (Figure 6, GFP probe, green asterisks). Insertion of more than one mEGFP-tag is also possible resulting in a shift of the tagged DNA fragment as demonstrated in clones #1 and #18 (Figure 6, GFP and NUP358 blot, clone #1 and #18, green asterisks). The expression of randomly inserted tags should be investigated thoroughly using Western blot analysis, since extra FP can interfere with the outcome of downstream experiments such as quantitative live cell imaging (see our accompanying Nature Protocol *et al.*^4^).

**Figure 6:**
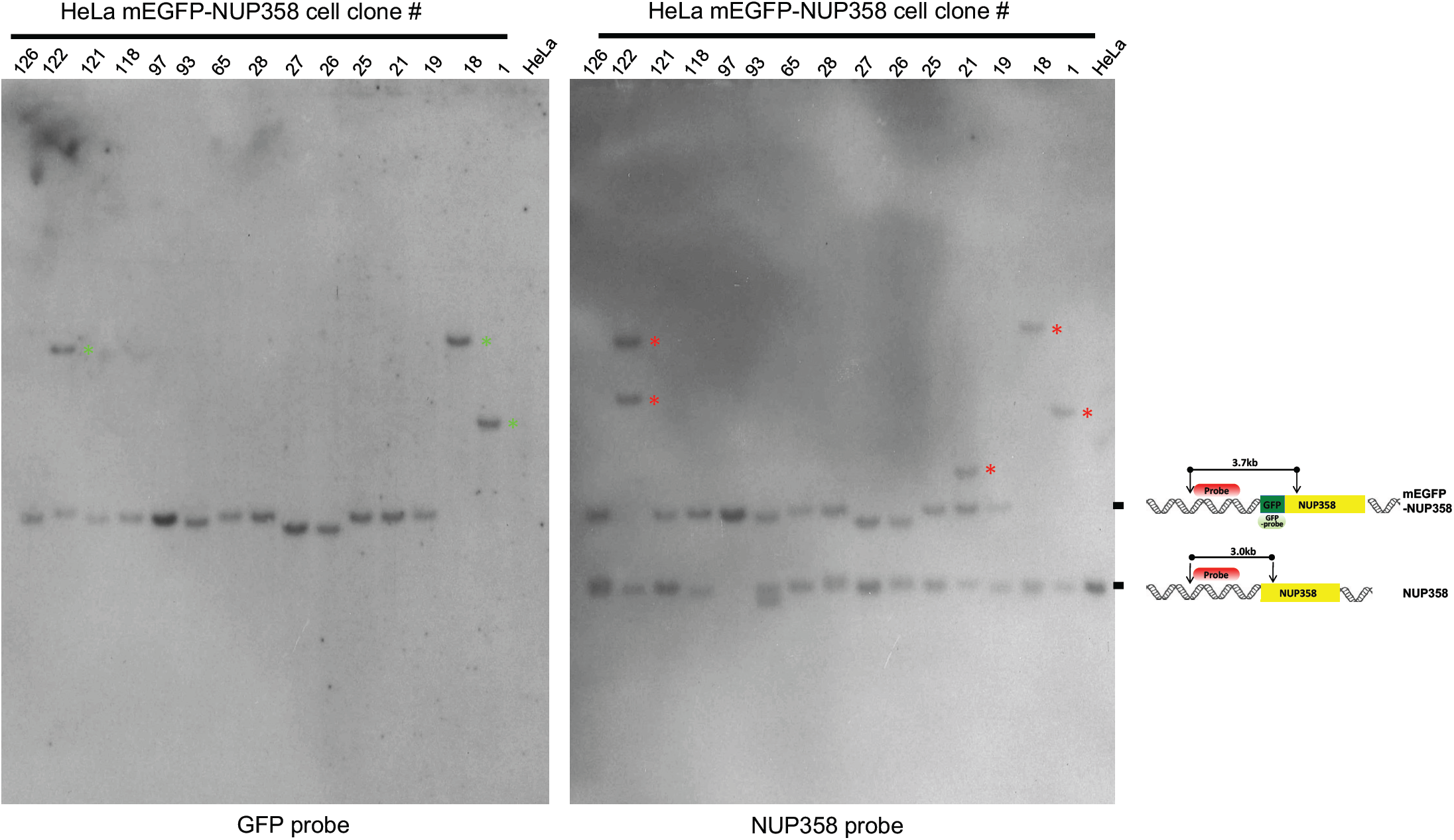
Southern blot analysis of HeLa Kyoto mEGFP-NUP358 cell clones. Southern blot was performed with different HeLa Kyoto cell clones expressing mEGFP-NUP358 using GFP or NUP358 probes as described in step 48 Option B. Briefly the genomic DNA was digested with BamHI and SphI. GFP probe is binding within the GFP gene and can be detected as a 3.7kb fragment. The NUP358 probe which was design to bind at the 5’ end of NUP358 outside the left homology arm can detect either the untagged NUP358 (= 3.0kb fragment) or tagged mEGFP-NUP358 (= 3.7kb fragment). The scheme on the right hand side depict the binding of the probes and the expected results. Homozygous clones will lead to the detection of only the tagged mEGFP-NUP358 (3.7kb fragment) as demonstrated with clone #97. Green asterisks in the blot with GFP probe indicate extra integration of mEGFP. Red asterisks in the blot with NUP358 probe reveal genomic rearrangements of the chromosome within this region. (*) = additional or shifted DNA fragments.

Probes binding to the endogenous locus of interest outside of the homology arms detect hetero- and homozygosity (Figure 6, NUP358 probe, Nup358 and mEGFP-NUP358). Homozygous clones result only in one expected product with the same size as for the probe against the tag (Figure 6, clone #97) whereas heterozygous clones have two products, one for the tagged gene and one for the untagged gene (Figure 6, e.g. clone #118). Rearrangements within the endogenous locus of interest can occur and are detected by extra unexpected bands (Figure 6, NUP358 probe, red asterisks). Furthermore some samples, such as #26 and #27, can show products with a lower size of the tagged GOI as expected which could be due to deletions within this region.

Taken together clones that show no extra unexpected DNA fragments or unexpected pattern or sizes are used for further experiments.

### Western blot analysis

Expression of the fluorescently tagged protein is further characterized by Western blot analysis (Figure 7). An antibody against the FP, such as anti-GFP, detects specific expression of the tagged proteins of interest as a specific band at the expected size (Figure 7, upper panel anti-GFP, mEGFP-NUP358 = 385 kD). Random integration of the donor plasmid can cause expression of free FP when it is inserted in euchromatin regions, which is detectable as a 27 kDa protein band. There is a weak unspecific band at the same size as free GFP which is visible in the HeLa control sample and which in turn could interfere with the detection of free GFP. This unspecific band is most likely due to unspecificity of the antibody which may be solved by using a different antibody. GFP antibodies from various companies, such as Abcam, Santa Cruz, Rockland, MBLand Roche, were tested and the one generated by Roche was the most sensitive and specific one so far. However, HeLa Kyoto mEGFP-NUP358 clones #26 and #122 seem to express low amounts of free mEGFP, as a 27 kDa band is detectable with the GFP antibody (Figure 7, upper panel, anti-GFP), which is elevated in comparison to HeLa Kyoto wt cells.

**Figure 7:**
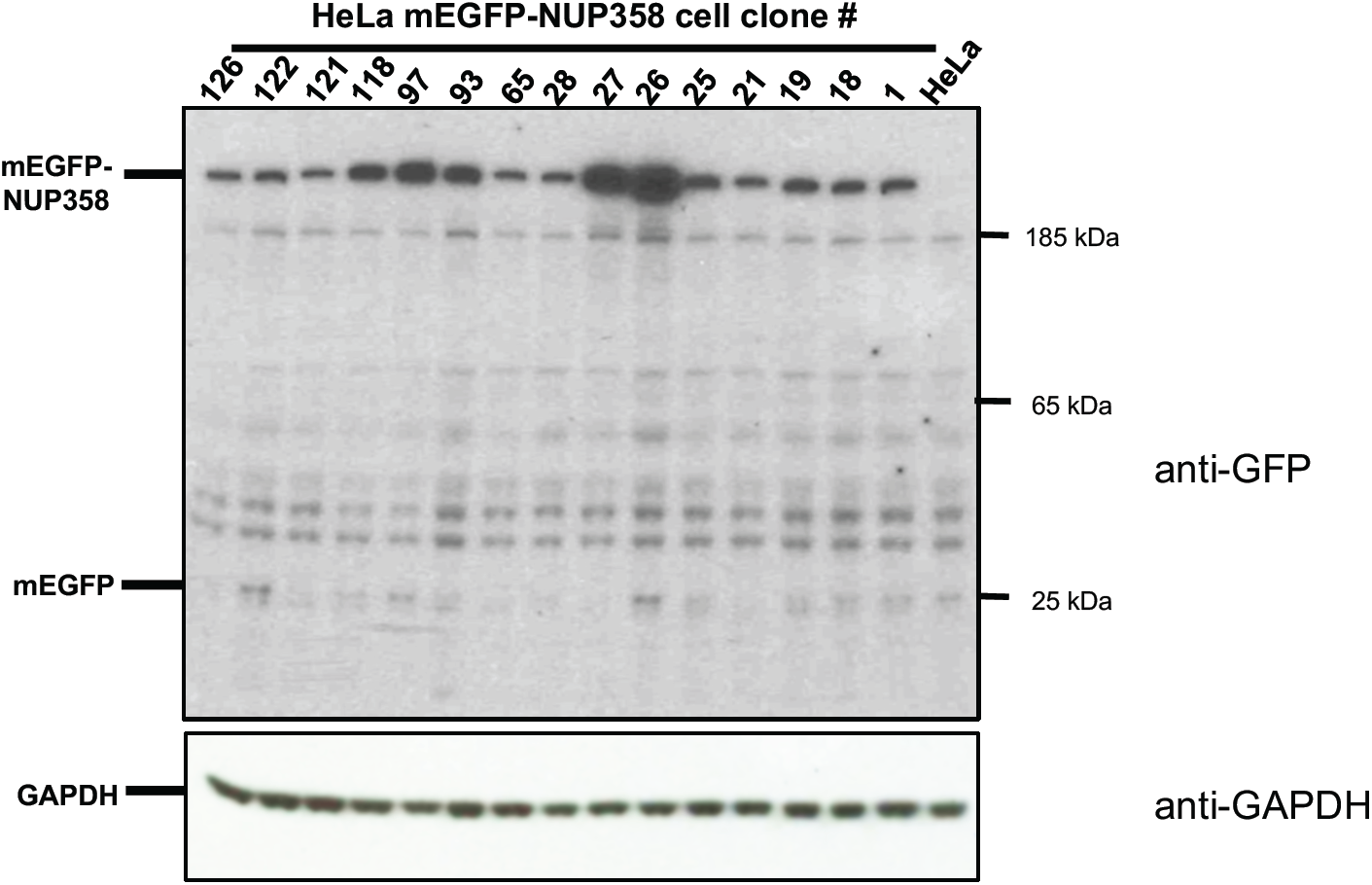
Western blot analysis of HeLa Kyoto mEGFP-NUP358. An antibody against GFP (upper panel, Roche cat no. 11814460001) was used to detect full length expression of the endogenously tagged mEGFP-NUP358 protein. Anti-GAPDH (lower panel, Santa Cruz cat no. sc-32233) was used as loading control.

Expression of free FP can interfere with protein characterization experiments such as quantitative live cell imaging. Consequently, clones expressing free GFP above the background levels, should not be used for further experiments. Similar to GFP, Western blot analysis with antibodies against the used tags, which were endogenously introduced, have been successfully used to detect expression of other free FPs/tags.

A specific antibody against the POI, if available, is very useful to distinguish homo- and heterozygous clones as shown in Mahen *et al*.^1^. However, in the case of mEGFP-NUP358 this is not feasible since the differences between the 358kD untagged NUP358 and 385kD tagged mEGFP-NUP358 cannot be detected by routine Western blot analysis due to the large size of the protein.

### Cell cycle analysis

The cell cycle can be influenced by the tag or off-target effects after the genome editing process. A major interest in our group is the characterization of proteins involved in cell division and in cell cycle control. We therefore established an automated workflow consisting of automated time-lapse microscopy and its analysis to measure the cell cycle kinetics of many cells at once (Figure 8). Similar assays for other cellular functions of interest can be established using high-throughput microscopy with appropriate markers ^30^.

**Figure 8:**
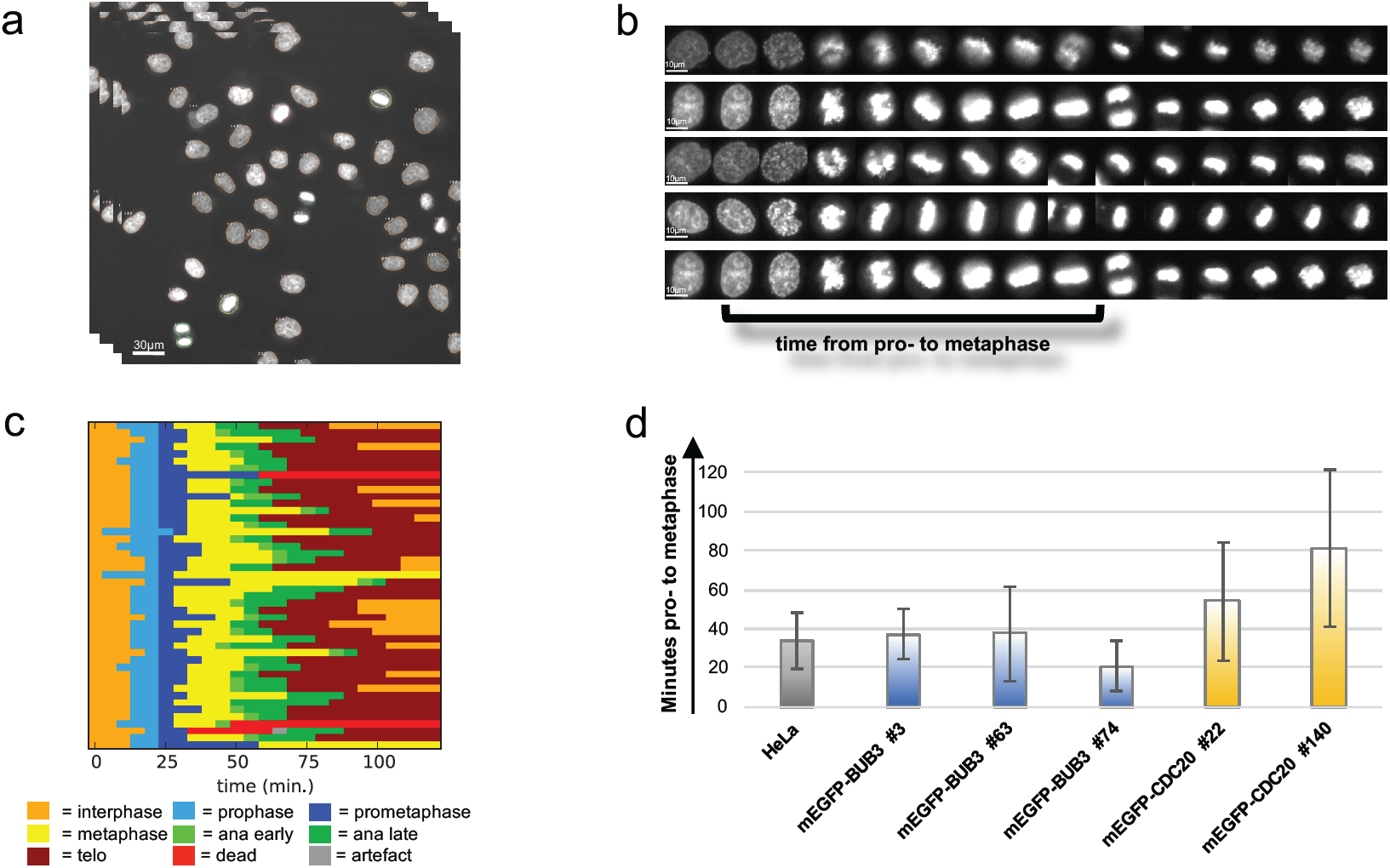
Workflow of the analysis of mitotic timing. (a) Automated live cell imaging of SiR-DNA stained cells to detect the timing of mitosis. Pictures were taken every 5 min. over 24h (scale bar = 30μm) (b) Live cell images were automatically analysed using CellCognition software to track single cells and classify them into interphase and mitotic phases (scale bar = 10μm). Subsequently, (c) the mitotic timing of each single cell was analysed. The different colors represent different mitotic phases for every single cell. (d) Representative statistical evaluation of the duration of mitosis for various cell lines expressing various tagged proteins. The error bars are the standard deviations of the mean mitotic timing from proto onset of anaphase.

First, the nuclei of the cells are stained with SiR-DNA to determine the cell cycle status of each individual cell. Subsequently automated live cell imaging is performed every 5 min for 24h (Figure 8A). The images are automatically analyzed to track single cells, detect interphase and mitotic cells, and measure mitotic timing (Figure 8B and C). For instance, the mitotic time from the start of prophase to the onset of anaphase is measured (Figure 8C) and statistically evaluated (Figure 8D). The duration of mitosis is compared to wildtype cells. Cell clones with significantly extended (Figure 8D, mEGFP-CDC20 #140) or shortened mitotic time (Figure 8D, mEGFP-Bub3 #74) are discarded.

The mitotic time of all cell clones per tagged gene are analyzed. For example, all clones of HeLa Kyoto mEGFP-NUP358 cells behave as HeLa Kyoto wt cells during mitosis (Figure 9), validating their use in future experiments, such as FCS-calibrated 4D live cell imaging ^2,4^.

**Figure 9:**
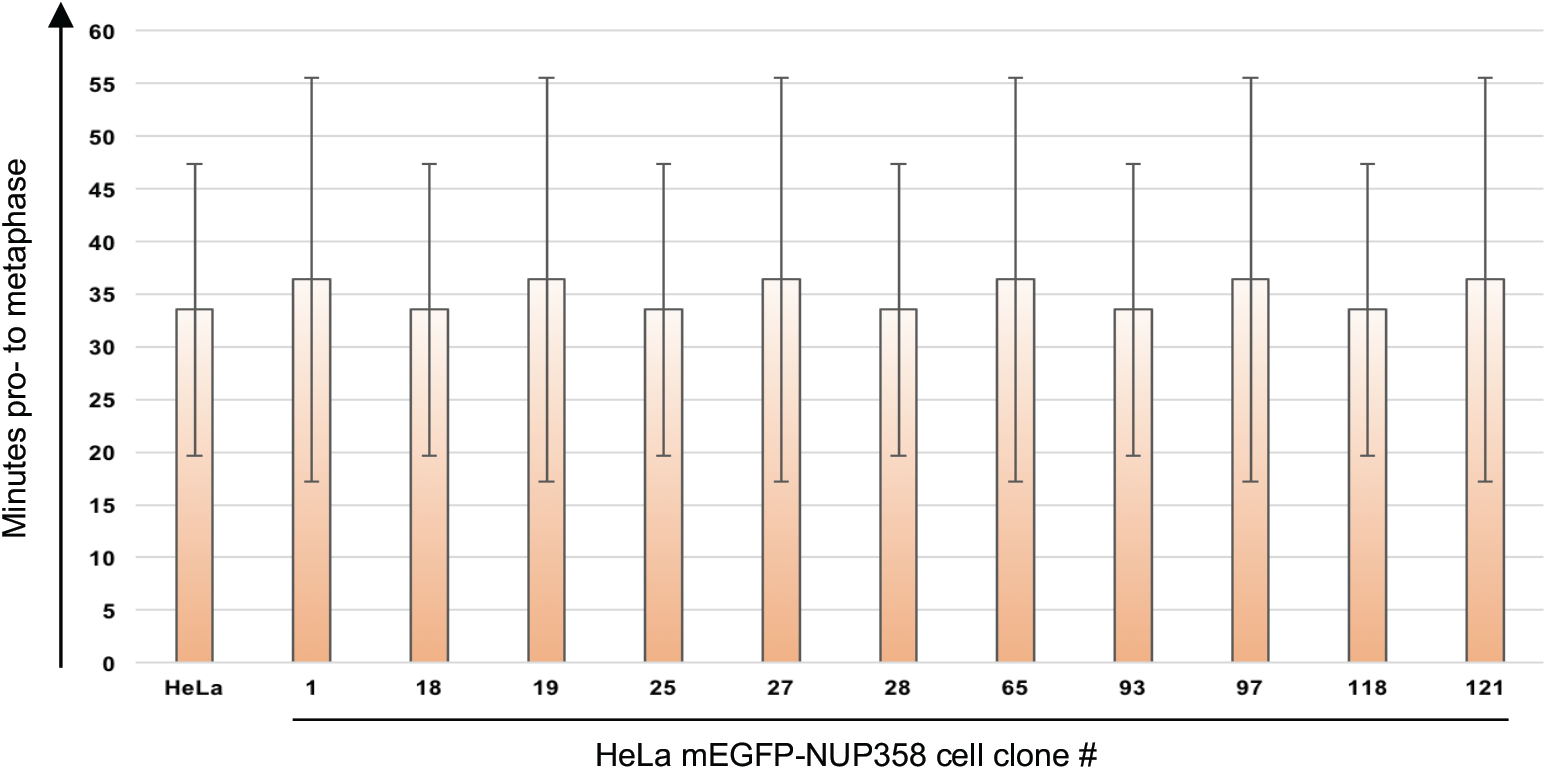
Mitotic duration of HeLa Kyoto mEGFP-NUP358 cell clones. Results of the mitotic analysis workflow for all HeLa Kyoto mEGFP-NUP358 cell clones. The duration of mitosis of HeLa Kyoto mEGFP-NUP358 clones was compared to HeLa Kyoto wt cells. The error bars are the standard deviations of the mean mitotic timing from pro-to onset of anaphase.

### Required controls

The advantage of endogenously tagging POIs is that tagged proteins keep not only physiological expression levels, but also keep their functionality. However, possible off-target effects and the FPs can influence these. Therefore, it is essential to develop a validation workflow to assay for correct expression and functionality of the POIs.

There are several controls to be considered during the generation and validation workflow. The controls are prepared exactly as the samples. The preparation of each sample/control at each validation step is described in detail in the procedure section, however here we present a summary of the required controls.

The negative control for the functionality test of gRNAs is the untransfected cells control and as positive control already validated gRNA pairs can be used (step 26). The negative controls should only have one band as the PCR product band whereas the paired nickase sample digested with T7E1 should have several cleavage bands as predicted. An alternative to the T7E1 assay is TIDE. TIDE consists of the amplification of the target region, Sanger sequencing and analysis of the sequence traces by a specially developed decomposition algorithm that identifies the major induced mutations in the projected editing site and accurately determines their frequency in a cell population ^43^.

The expression levels of endogenously tagged protein during FACS can be very low. Therefore, it is essential to use HeLa Kyoto wt cells as negative control to distinguish between non-specific and specific FP/mEGFP expression levels (step 42).

The genomic DNA from parental cell lines such as HeLa Kyoto and water can be used as negative controls to validate the integration of the tag at the correct locus and test of homozygosity by junction PCR (step 48 Option A).

Genomic DNA prepared from parental cell lines such as HeLa Kyoto cells can be used as negative controls of Southern blot analysis (step 48 Option B) and Sanger sequencing (step 48 Option C) to distinguish between tagged and untagged DNA.

In Western bot analysis protein extracts derived from parental cells can be used to compare expression levels of untagged versus tagged protein (step 48 Option D).

The correct localization of the tagged POI can be confirmed by the use of antibodies against the POI during microscopy of fixed cells ((step 49 Option B).

The cell cycle of the endogenously tagged cell lines are compared to the cell cycle timing of the parental cells such as HeLa Kyoto cells (Box 1).

Taken together, the expression and function of the POI in the parental cell line in comparison to the endogenously tagged POI is essential. Although if the POI is unknown, it will be difficult to evaluate these and most likely several tools such as antibodies have to be additionally generated.

## MATERIALS

### REAGENTS

#### gRNA cloning

- pX335-U6-Chimeric_BB-CBh-hSpCas9n(D10A) (Addgene plasmid ID#42335)
- Oligos for gRNA construction (see Table 1)
- NEB^®^ 5-alpha Competent *E. coli* (High Efficiency) (New England Biolabs, cat. no. C2987H)
- QIAquick gel extraction kit (QIAGEN, cat. no. 28704)
- QIAprep spin miniprep kit (QIAGEN, cat. no. 27106)
- SYBR Safe DNA stain, 10,000× (Life Technologies, cat. no. S33102)
- FastDigest BbsI (BpiI) (Thermo Scientific, cat. no. FD1014)
- Roche Rapid Ligation Kit (Roche, cat. 11635379001)
- NaCl (Merck, cat. S7653)
- Peptone special (Merck, cat. 68971)
- Yeast (Merck, cat. 51475)
- Agar (merck, cat. A5306)
- Ampicillin (Sigma, cat. no. A9518)
- Agarose (Sigma, cat. no. A9539)
- 10X TBE (ThermoFisher, cat. AM9864)
- Agarose (Merck, cat. A9539)
- Gene ™ Ruler DNA Ladder Mix (Thermo Scientific, cat. no.SM0331)
- Loading Dye (6x) (Thermo Scientific, cat. no. R0611)

#### T7E1 assay

- HotStar HighFidelity PCR kit (QIAGEN, cat. no. 202605)
- Oligos
- T7 Endonuclease I (New England BioLabs, cat. no.M0302S)
- NEB buffer 2 (New England BioLabs, cat. no. B7002S)

#### Mammalian cell culture

HeLa Kyoto cells (from S. Narumiya, Kyoto University Biosafety level S1). The cells are not known to be misidentified no cross-contaminated. The cell lines are regularly checked and not infected with mycoplasma.

CAUTION: The cell lines used in your research should be regularly checked to ensure they are authentic and are not infected with mycoplasma.

- High Glucose Dulbecco’s Modified Eagle Medium (DMEM) (Life Technologies, cat. no. 41965-039)
- Fetal Bovine Serum (FBS, Life Technologies, cat. no.10270106)
- 10000 units/ml Penicillin – Streptomycin (Life Technologies, cat. no. 15140122)
- 200 mM L-Glutamine (Life Technologies, cat. no. 25030081)
- KCl (Merck, cat. P9333)
- Na_2_HPO_4_ (Merck, cat. S7907)
- KH_2_PO_4_ (Merck, cat. P5655)
- 0.05% Trypsin (Life Technologies, cat. no. 25300054)

#### Transfection

- Donor plasmids synthesized from Geneart (Thermo Scientific) or Genewiz (Sigma Aldrich)
- jetPrime (Polyplus Transfection, cat.no. 114-07)
- Opti-MEM^®^ I Reduced Serum Medium, GlutaMAX™ Supplement (Life Technologies, cat.no. 51985-026)

#### FACS

- BD FACSFlow ™ (Becton Dickinson GmbH, Cat. No. 322003)
- Flow-Check ™ Fluorospheres (Beckman Coulter, Cat No. 6605359)

#### Junction PCR

- HotStar HighFidelity PCR kit (QIAGEN, cat. no. 202605)
- Trizma^^®^^base (Sigma, cat.no. T1503)
- TWEEN^®^ 20 (Merck, cat. P1379)
- Triton X100 (Merck, cat. T9284)
- MgCl_2_ (Merck, cat. M8266)
- Proteinase K (Merck, cat. P2308)

#### Southern blot analysis

- PCR DIG probe Synthesis kit (Roche, cat. no. 11 636 090 910)
- Blocking agent (Roche, cat. no. 1109617600)
- Epicentre MasterPure DNA purification kit (cat. no. MCD85201)
- DIG Easy Hyb Buffer (Roche, cat. no. 11603558001)
- anti-digoxigenin-Alkaline Phosphatase Fab fragments (Roche, cat. no.11093274910)
- CDPstar chemiluminescent substrate (Sigma, cat. no. C0712)
- High performance autoradiography film (GE Healthcare, cat. no. 28906845)
- Sodium citrate (Merck, cat. 71497
- SDS (Merck, cat 75746) **! CAUTION** This compound is harmful. Avoid inhalation and contact with skin and eyes. Refer to the MSDS before use.
- NaOH **! CAUTION** NaOH is highly corrosive, causing chemical burns and irritation of the respiratory system. Avoid inhalation or contact with skin and eyes, and refer to the MSDS before use.
- Maleic Acid (Merck, cat. M0375)
- Zero Blunt ^®^TOPO^®^PCR Cloning Kit (Life Technology, cat. no. K2800-20)
- Kodak BioMax MR films (Sigma Aldrich, cat. no. Z350370-50EA)

#### Sanger Sequencing

- Oligos for amplification of target sequence and sequencing (see Table 1)
- HotStar HighFidelity PCR kit (QIAGEN, cat. no. 202605)
- MinElute Gel Extraction Kit (QIAGEN, cat. no. 28604)

#### Western blot analysis

- Cas9-HRP antibody (abcam, cat. no. ab202580)
- GFP antibody (Roche, cat. no. 11814460001)
- RFP antibody [3F5] (ChromoTek, cat. no. 3f5)
- Non-Fat Dry Milk powder
- DTT (Merck, cat. D0632)
- cOmplete EDTA-free Protease Inhibitor Tablets (Roche, cat. no.11873580001)
- PhosStop Tablets (Roche, cat.no. 04906845001)
- Glycerol (Merck, cat. G5516)
- Bio-Rad Protein Assay (Bio-Rad, cat no. 500-0006)
- NuPAGE^^®^^4-12%Bis-Tris Gels (Thermo Scientific, cat. no. NP0323BOX)**! CAUTION** Gels contain Acrylamide which is toxic and gloves should be worn.
- Western LIGHTNING ™ *plus*-(Perkin Elmer, cat. no. NEL105001EA) or Amersham ™ ECL ™ Prime Western Blotting Detection Reagent (GE Healthcare, cat. no. RPN2232)
- NuPAGE MOPS SDS Running Buffer, 20X (Thermo Scientific, cat. no. NP0001)
- NuPAGE LDS sample buffer, 4X (Thermo Scientific, cat. no. NP0008)
- PageRuler Plus Prestained Protein Ladder 10-250 kDA (Thermo Scientific, cat no. 26619)
- Immobilon-P Transfer Membrane (PVDF membrane, Merck, cat. no. IPVH00010)
- 3MM Whatman paper

#### Live cell imaging and cell cycle analysis

- SiR-DNA (SPIROCHROME, cat. no. SC007)
- CO_2_-independent imaging medium without phenol red (Gibco, cat. no. 18045088 custom made without phenol red)
- Sodium pyruvate (Gibco, cat. no. 11360070)
- Silicon grease (Bayer, cat. ACRO386110010)
- 16% PFA (ThermoFisher, cat. 28906) **! CAUTION** Concentrated formaldehyde is toxic, and should be handled under a hood.
- Hoechst 33342 (ThermoFisher, cat. no. H3570)

## EQUIPMENT

- NuncDish 24 well (Thermo Scientific, cat. no. 142475)
- NuncDish 6 well (Thermo Scientific, cat. no. 140675)
- NuncDish 96well (Thermo Scientific, cat. no. 167008)
- Cryotube™ vials Thermo nunc (Thermo Scientific, cat. no. 366656)
- DAKO MoFlo High Speed sorter
- Falcon^®^ 12 x 75 mm Tube with Cell Strainer Cap (Falcon, cat.no. 352235)
- Falcon^®^ 12 x 75 mm Tube Snap Cap (Falcon, cat.no. 352058)
- Falcon 14 ml polypropylene tubes (Thermo Scientific, cat.no. 352059)
- CellStar tubes 15 ml, PP, conical bottom (Greiner, cat. no. 188271)
- Falcon^™^ Disposable Polystyrene Serological 5 ml Pipets (Falcon, cat. no. 357543)
- Falcon™ Disposable Polystyrene Serological 10 ml Pipets (Falcon, cat. no. 357551)
- Falcon^™^ Disposable Polystyrene Serological 25 ml Pipets (Falcon, cat. no. 357525)
- Filter tips: 1000 μl, 200 μl, 20 μl, 10 μl
- 3MM Whatman paper
- Genescreen Plus Hybridization Transfer Membrane (Perkin Elmer,cat. no. NEF976001PK)
- Immobilon-P Transfer Membrane (PVDF membrane, Merck, cat. no. IPVH00010)
- 96 Well SCREENSTAR Microplate (Greiner, cat. no. 655866)
- Imaging plates 96CG, glass bottom (zell-kontakt, cat. no. 5241-20)
- Glass hybridization flasks 40×300mm
- 0.22μm Filters
- Disposable cuvettes made of PS, semi-micro, 1.5-3.0 ml (neoLab, cat. no. E-1642)
- REDTM Imaging System (Biozym Scientific GmbH)
- Mini Trans-Blot^®^Electrophoretic Transfer Cell (BioRad, cat no. 1703930)
- ImageXpress Micro XLS Widefield High-Content Analysis System (Molecular Devices)
- PowerPac™ Basic Power Supply (BioRad, cat. no.164-5050)
- Rocker or shaker
- Hybaid Oven Mini10
- Kodak RPX-OMAT processor
- EI9001-XCELL II Mini Cell (NOVEX) gel chamber
- Transferpette-8 electronic 1-20 μl (Brand, cat. no. 705400)
- Transferpette-8 electronic 20-200 μl (Brand, cat. no. 705404)
- Submerged Horizontal agarose gel electrophoresis system

## REAGENTS SET UP

### Agarose gel

Resuspend the correct amount (%w/v) of agarose in 1X TBE. Boil until agarose is dissolved and cool down to ∼50-60°C. Dilute SYBR^®^ Safe DNA Gel Stain 100,000 fold in the gel prior to polymerization. Prepare freshly.

### Cell lysis buffer

Prepare a solution containing 10% glycerol, 1 mM DTT, 0.15m EDTA, 0.5% TritonX100 in 1x PBS. Dissolve 1 tablet of cOmplete EDTA-free protease inhibitor and 1 tablet of PhosStop in 10 ml cell lysis buffer. Prepare buffer freshly prior use.

### Denaturation Solution

Dissolve NaOH pellets to a final concentration of 0.5M and NaCl to 1.5M in ddH_2_O. Prepare freshly.

### DIG1, 10X

Dissolve Maleic Acid to a final concentration of 1M and NaCl to 1.5M in ddH_2_O. Adjust pH to 7.5 with NaOH pellets. Careful Maleic acid only dissolves at a pH above ∼6.5-7 and has hardly any buffering capacity. Filter sterilize and store at room temperature (RT, 23-25°C) up to 2 month.

### DIG1, 1X

Dilute 10X DIG1 buffer to 1X with ddH_2_O. Store at RT up to 2 months.

### DIG1 wash buffer

Dilute 10X DIG1 buffer to 1X with ddH_2_O and add 0.3% TWEEN^®^ 20. Store at RT up to 2 months.

### DIG3

Dilute stock solutions to final concentrations of 0.1M NaCl and 0.1M Tris (pH9.5) in ddH_2_O. Store at RT up to 2 months.

### DIG block, 10X

Weigh out 10% w/v Roche blocking agent and dissolve in 1X DIG1. Heat the solution to ∼65°C bringing the blocking reagent into solution. Make 10 ml aliquots and store at −20°C up to ∼ 6 months. Use a fresh aliquot every time due to the risk of contamination.

### DNA lysis buffer

Dilute and dissolve the following reagents in ddH_2_O: 0.45% TWEEN^®^ 20, 0.45% Triton X100, 2.5 mM MgCl_2_, 50 mM KCl, 10 mM Tris-HCl pH8.3. Store at RT up to 1 month and add 100ug/ml proteinase K prior to use.

### DTT, 1M

Prepare the solution in ddH2O and divide into 100 μl aliquots. Store them at −20°C up to 6 months. Use a new aliquot for each reaction, as DTT is easily oxidized.

### FACS buffer

Dilute 2% FBS in PBS, filter through a 0.22μm membrane. Prepare buffer freshly prior to use.

### HeLa growth medium

High Glucose DMEM supplemented with 10% FBS, 100 U/ml Penicillin, 100 μg/ml Streptomycin and 2 mM L-Glutamine. Supplemented DMEM medium is abbreviated as complete DMEM. Store at 4°C up to 2 weeks.

### Imaging medium

CO_2_ independent medium without phenol red supplemented with 20% FBS, 2 mM L-Glutamine and 1 mM sodium pyruvate. Filter through a 0.22 μM membrane to clear the medium of precipitants. Store at 4°C up to 2 weeks.

### LB medium

Dissolve 1% NaCl, 1% peptone and 0.5% yeast extract in ddH_2_O. Sterilize by autoclaving. Store at 4°C up to 2 months.

### LB agar medium

Dissolve 15 g Agar in 1l LB medium. Sterilize by autoclaving. Store at 4°C up to 2 months.

### NaCl, 5 M

Dissolve the appropriate amount of NaCl in ddH_2_O. Filter sterilize or autoclave. Store at RT up to 6 months.

### Neutralization solution

Dissolve NaCl to a final concentration of 1.5 M and Tris to 0.5 M in ddH_2_O. Adjust pH to 7.0 using HCl. Prepare freshly prior to use.

### PBS, 1X

Prepare 137 mM NaCl, 2.7 mM KCl, 10 mM Na_2_HPO_4_ and 1.8 mM KH_2_PO_4_ in ddH_2_O. Adjust the pH to 7.4 with HCl and autoclave. Store at RT up to 6 months.

### PBS-T

Add 0.1% TWEEN^®^ 20 into 1X PBS. Store at RT up to 2 months.

### SDS PAGE running buffer

Dilute 20X NuPAGE MOPS SDS Running Buffer to 1X with ddH_2_O. Prepare freshly prior to use.

### SDS sample buffer

4X NuPAGE^®^ LDS Sample Buffer was supplemented with 100 μM DTT and diluted with whole protein extracts to 1X. Prepare freshly prior to use.

### 20%SDS

Weigh out 20% w/v SDS pellets and dissolve in the appropriate amount of ddH_2_O by heating to 65-68°C and sterile filter the solution. Store at RT up to 6 months.

### SSC, 20X

Dissolve NaCl to a final concentration of 3 M and sodium citrate to 0.3 M in ddH_2_O. Adjust pH to 7.0 with 1 M HCl due to weak buffering capacity. Store at RT up to 6 months.

### Stripping solution

Dissolve NaOH pellets to a final concentration of 0.2 M NaOH in ddH_2_O and add 0.1% SDS. Prepare freshly.

### TBE, 1X

Dilute a 10X stock solution with ddH_2_O. The final concentration of 1X TBE is 89 mM Tris pH 7.6, 89 mM boric acid, 2 mM EDTA. Store at RT up to 6 months.

### 1M Tris pH9.5

Dissolve Trizma base to a final concentration of 1M in ddH_2_O and adjust the pH to 9.5 with concentrated HCl. Filter sterilize or autoclave. Store at RT up to 6 months.

### 1M Tris pH8.3

Dissolve Trizma base to the final concentration of 1M in ddH_2_O and adjust pH to 8.3 with concentrated HCl. Filter sterilize or autoclave. Store at RT up to 6 months.

### Western transfer buffer

Dilute NuPAGE^®^ (20X) Transfer Buffer in ddH_2_O to 1X and add 10% pure Ethanol. Prepare freshly prior to use.

### Western blot blocking solution

Resuspend 5% w/v non-fat dry milk powder in PBS-T. Store at 4°C for up to a few days.

## PROCEDURE

### gRNA design TIMING 1 d

**1.** *gRNA design.* Design a pair of gRNA, one binding to the antisense strand and the other to the sense strand, so that one of them binds on or as close as possible to the target site, e.g. ATG or Stop codon. Homologous recombination will occur most efficiently within 100 bp of the target site and drops dramatically beyond this region. Design two different gRNA pairs using either http://crispr.mit.edu/ or https://benchling.com/crispr ^9, 44, 45^. Choose gRNAs with predicted off-targets that contain at least 2nt mismatches located within the seeding region 8-12 nt 5’of the PAM sequence.

**2.** *Design of oligos for gRNA cloning.* Digest the pX335 vector using BbsI, and clone a pair of annealed oligos into the pX335 vector before the gRNA scaffold. Design oligos based on the target site sequence (20 bp) where the 3bp NGG PAM sequence is at the 3’ end but do not include the PAM site. BbsI sites were added to the gRNAs to enable cloning the oligos into pX335 vectors as described on the webpage https://www.addgene.org/crispr/zhang/ (see Table 1).

**3.** Order the oligos (shown in Table 1).

### Donor design and synthesis TIMING 14-21 d

**4.** *Donor design* Design the donor that the plasmid DNA contains the fluorescent marker gene (e.g. mEGFP or mCherry) which is flanked by 500-800 bp homology arms of the GOI. The fluorescent marker replaces either the start or the stop codon and is in-frame with the GOI and the endogenous promoter. If the ends of the homology arms are GC rich or contain any repetitive regions, they should be shortened, otherwise 800 bp works well. A linker should be placed between the tag and the gene to maintain functionality of the tagged protein. You can use 5xGlycine or 5x Glycine/Serine repeats as a linker ^46^ however, if there is a known linker that has already been used for the POI, it is best to use that one. The inclusion of additional cloning sites between the homology arms and the insert enables the exchange of the tag or linker if required (Supplementary Figure S1). If two proteins of interest have to be tagged, the least expressed protein should be tagged with mEGFP since it is brighter and more stable than mCherry. Use monomeric fluorescent tags such as mEGFP, mCherry or mCer3. The donor design usually depends on the gRNAs as it should differ from the endogenous gene otherwise the gRNA will also cut the donor, causing a decrease in HDR efficiency. Therefore, it is best to use gRNAs across the start or stop codon (depending on N-or C-terminal tagging, respectively), so that the donor differs to the endogenous genomic locus due to the insertion of the tag.

**CRITICAL STEP** No selection marker for mammalian cells is contained in the donor plasmid. If antibiotic selection in mammalian cells is used, random integration of the donor plasmid into the cell genome will increase. Also, no CMV, RSV or any other exogenous promoter is present within the donor plasmid, otherwise the expression levels of the tagged protein will not be physiological and artefacts will occur.

**5.** *Gene synthesis of donor* Synthesize the donor using either Geneart (Thermo Scientific) or Genewiz (Sigma Aldrich).

**CRITICAL STEP** The synthesis of the donors can take up to 21 days depending on the G/C % and repetitive sequences in the homology arms. During the production period, proceed with the gRNA cloning and the T7E1 assay.

### Cloning gRNAs into the pX335-U6-Chimeric-BB-CBh-hSpCas9n(D10A) vector TIMING 4 d

**6.** Resuspend the forward and reverse strand for each gRNA oligo to a final concentration of 100μM in ddH_2_O.

**7.** Use 4 μl of each 100μM stock solution of forward and reverse oligo and mix well by pipetting.

**8.** Anneal forward and reverse oligo (Table 1) in a thermocycler by using the following parameters: 95°C for 5min. and subsequently cool to 25°C with a ramp of 0.1°C/sec (∼ 5°C/min.)

**9.** Combine the following components to digest pX335-U6-Chimeric-BB-CBh-hSpCa9n(D10A) vector with BbsI (BpiI):

| Component | Amount | Final concentration |
| --- | --- | --- |
| pX335-U6-Chimeric-BB-CBh-hSpCas9n(D10A) vector | 2 μg | |
| FastDigest BbsI (BpiI) | 2 μl | |
| 10X Fast Digest Buffer | 4 μl | 1X |
| ddH<sub>2</sub>O | up to 40 μl | |

Incubate digestion mix at 37°C for 30min.

**10.** After the incubation period, add 8μl of 6X loading dye to 40μl of the digestion product, load on a 1% (w/v) agarose gel in TBE buffer with SYBR Safe dye and run at 100V for ∼ 20min.

**11.** Purify the digestion product of 8.4kb using the QIAquick gel purification kit according to the supplier’s manual. Elute the DNA in 30 μl of EB buffer (part of the kit).

**12.** *Ligation of annealed oligos into BbsI-digested pX335 using the Roche Rapid Ligation Kit.* Prepare for each gRNA and a no-insert negative control the following ligation reaction:

| Component | Amount |
| --- | --- |
| BbsI-digested pX335 vector | 50-100 ng |
| Annealed gRNA oligos | 1 μl (replace with water for no-insert control) |
| 5X DNA dilution buffer | Up to 10 μl |
| T4 DNA ligation buffer | 10 μl |
| T4 DNA ligase | 1 μl |

Incubate the ligation mix at RT for more than 5min.

**CRITICAL STEP** In former protocols^9^, phosphorylation of the annealed oligos and dephosphorylation of the vector were performed, which may cause an elevated number of false positive colonies or no colonies at all. The above protocol is reliably produces positive colonies for the correct insertion of gRNA oligos, although you may find that sometimes only few colonies are generated.

**13.** *Transformation of ligation reactions into a competent E.coli strain.* Add 10 μl of each ligation reaction into 50 μl ice-cold chemically competent NEB 5alpha competent bacteria, incubate on ice for 30min, perform a heat-shock at 42°C for 45sec and place on ice immediately for 2-3min.

**14.** Add 1 ml RT SOC medium to the mixture and incubate at 37°C with agitation for 60 min. Subsequently spin at RT for 1min. at 3000g, remove supernatant and resuspend bacteria pellet in 100 μl LB medium.

**15.** Plate the suspension on a LB agar plate containing 100 μg/ml ampicillin and incubate at 37°C overnight.

**16.** Day 2: Examine the LB plates and pick a few colonies, inoculate these into 5 ml LB medium containing 100 μg/ml ampicillin. Incubate these cultures at 37°C overnight.

**17.** Day 3: Isolate the plasmid DNA from the LB cultures by using QIAprep spin miniprep kit according to the supplier’s manual.

**18.** Validate the insertion of the gRNA into pX335 plasmids (30-100ng/μl) by Sanger sequencing ^41,42^ using the following oligo: 5’ TGCATATACGATACAAGGCTGTTAG 3’ (Table 1). Sanger Sequencing services are available from various companies such as GATC and details for the preparation of DNA for sequencing are found for example on the GATC home page http://www.gatc-biotech.com/en/index.html). Permanently store the plasmids at −20°C.

**TROUBLESHOOTING**

### Functionality test of gRNAs using the T7E1 assay TIMING 5 d

**19.** *Maintenance of HeLa Kyoto cells. Culture* HeLa Kyoto cells in High Glucose DMEM medium supplemented with 10% FBS, 1000 unit/ml PenStrep, 2 mM glutamine, 1 mM sodium pyruvate at 37 °C and 5% CO_2_. Cells were cultured in 10 cm cell culture dishes for up to 20 passages. Split cells at ∼ 80% confluency as described in steps 20-22.

**20.** Remove the medium and rinse the cells once with 1X PBS.

**21.** Add 1-2 ml of trypsin onto a 10 cm dish and incubate at RT until cells are almost detached, remove trypsin and resuspend cells in 5-10 ml of growth medium.

**22.** Transfer 1:5-1:10 of the cell suspension into a new 10 cm cell culture dish and put them back into an incubator with 37°C, 5% CO_2_.

PAUSE POINT Grow the cells until further use and split them at ∼ 80% confluency.

**23.** *Transfection of HeLa Kyoto cells with gRNA plasmids.* Seed 1.5×10^5^ cells per well on a 6 well cell culture dish in 2 ml of normal growth medium and incubate at 37°C, 5% CO_2_ for one day prior to the transfection. The cells should be at least 50% confluency during the transfection.

**24.** The next day dilute the gRNA plasmids in 200 μl of jetPrime buffer at the concentration indicated in the table below.

| Sample | CRISPR/Cas9<br>nickase sense | CRISPR/Cas9<br>nickase antisense | Buffer | JetPrime |
| --- | --- | --- | --- | --- |
| CRISPR Oligo 1<br>(sense)–negative<br>control | 2 µg |  | 200 µl | 4 µl |
| CRISPR Oligo 2<br>(antisense)–negative<br>control |  | 2 µg | 200 µl | 4 µl |
| Paired CRISPRs | 1 µg | 1 µg | 200 µl | 4 µl |
| Untransfected cells-<br>negative control | - | - | - | - |

**25.** Mix well and add 4 μl of jetPrime solution, incubate at RT for 15min.

**26.** Add transfection complex onto the cells from step 23.

**27.** Incubate 4h at 37°C and 5% CO_2._

**28.** Replace transfection medium with normal growth medium.

**29.** Incubate at 37°C and 5% CO_2_ overnight.

**30.** Day 2: Passage cells if they are more than 80% confluent.

**31.** Day 3: *Harvest cells for genomic DNA extraction 72h post transfection*. Wash cells with 1X PBS, add 1 ml trypsin and wait until cells almost detach. Remove trypsin and resuspend cells in 1 ml growth medium. Transfer cell suspension into an Eppendorf tube, spin at RT for 1 min. at 200g, wash cell pellet in 1X PBS and spin again for 1min. at 200g.

**32.** Prepare genomic DNA from cell pellets using the GenElute Mammalian Genomic DNA Miniprep Kit (Sigma) or a similar genomic DNA preparation kit.

**33.** *Touchdown PCR.* Design primers resulting in a final 700-1000 bp PCR product. Successful cutting of this PCR product via CRISPR/Cas9 should result in two fragments. Each of the cut fragments should be around 300 bp and differ 100-200 bp in length to each other resulting in two detectable bands on an 2% agarose gel. Amplify the region of interest using HotStar HighFidelity PCR kit and set up each reaction as described below:

| Component | Volume |
| --- | --- |
| 5x HotStar PCR buffer | 4 µl |
| T7 Primer_fw (10 µM) | 2 µl |
| T7 Primer_rev (10 µM) | 2 µl |
| HotStar HighFidelity polymerase | 1 µl |
| Q Solution | 4 µl |
| Template DNA (200 ng) |  |
| ddH<sub>2</sub>O | Up to 20 µl |

Amplify the region of interest using the following cycling conditions:

| Cycles | Denature | Anneal | Extend |
| --- | --- | --- | --- |
| 1 | 95 °C, 5 min |  |  |
| 2-21 | 95 °C, 30s | Start temperature is 5°C higher than T<sub>m</sub> of oligos, - 0.5°C/cycle, 45s | 72°C, 0.5kb/1min. |
| 22-43 | 95 °C, 30s | End temperature of last cycle = 5°C lower than T<sub>m</sub>, 45s | 72°C, 0.5kb/1min. |

**34.** Run 2 μl of PCR product on a 2% agarose gel. One single amplified fragment should be present. Keep 3 μl of PCR product on ice for later analysis in step 37.

**35.** Use the remaining 15 μl of PCR product and perform the following denaturing/reannealing cycle to generate heteroduplexed DNA:

| Temperature | Time |
| --- | --- |
| 95 °C | 5 min |
| 95 °C – 85 °C | -2 °C/s |
| 85 °C – 25 °C | -0.1 °C/s |
| 25 °C | 10 min |

**36.** Digest 15 μl of heteroduplexed DNA by adding 1.8 μl of 10X NEB buffer 2 and 1.2 μl of T7 Endonuclease I. Incubate at 37°C for 30min.

**37.** Analyze 18 μl of T7E1 digested product on a 2% agarose gel. Run 3 μl of undigested PCR product to compare the cleavage efficiency. The negative controls should only have one band as the PCR product band whereas the paired nickase sample digested with T7E1 should have several cleavage bands as predicted. The gRNA pair with the highest cleavage efficiency will be used in step 39 to introduce the fluorescent marker gene at the locus of interest.

**TROUBLESHOOTING**

### Delivery of paired CRISPR/Cas9 approach using plasmids TIMING 7-10d

**CRITICAL** The paired CRISPR/Cas9 approach has been used in a number of mammalian cell lines^6,7,16–18^. The transfection conditions for each cell line must be optimised. The conditions described below are best for HeLa Kyoto cells.

**38.** *Transfection of HeLa Kyoto cells with gRNA and donor plasmids.* Seed 1.5×10^5^ cells per well on a 6 well cell culture dish in 2 ml OptimemI medium to starve the cells prior transfection. Incubate the cells at 37°C, 5% CO_2_ for one day prior to the transfection. The cells should be at least 50% confluency during the transfection.

**CRITICAL STEP** Homozygosity can be increased up to 2% when cells are seeded into medium without serum such as OptimemI causing starvation and G1 arrest. Consequently, the cells enter S-phase after the transfection. Homologous recombination occurs more frequently during this stage of the cell cycle.

**39.** Day 2: Ensure the cells are at least 60% confluent during transfection. Dilute the chosen gRNA plasmids from step 37 and donor plasmid in 200 μl of jetPrime buffer as indicated in the table below. Mix well and add 4 μl of jetPrime solution, incubate at RT for 15min and add transfection complex onto the cells. After 4h at 37°C and 5% CO_2_ incubation replace transfection medium with normal growth medium and grow cells until they are∼80% confluent.

| CRISPR/Cas9<br>nickase sense | CRISPR/Cas9<br>nickase antisense | Donor<br>plasmid | Buffer | JetPrime |
| --- | --- | --- | --- | --- |
| 0.5 µg | 0.5 µg | 1 µg | 200 µl | 4 µl |

**40.** Day 3-10: Passage cells every time that they are reaching ∼80% confluency The single sort is performed with 1-2×10^7^ cells 7-10 days post transfection.

### Selection of single cell clones expressing fluorescent tag using FACS TIMING 10-12d

**CRITICAL** Single cell sorting can cause growth problems for some cell lines therefore it is important to establish the best growth conditions such as conditioned medium. The conditioned growth medium contains growth factors produced by the parental cell lines. Conditioned medium is a mixture of 1volume of medium in which the parental cell lines were grown for a few days and 1 volume of fresh complete growth medium. HeLa Kyoto cells survive single cell sorting relatively well and the survival rate is 30-50%. Perform FACS with at least one 10 cm cell dish (if possible with 1-2x 15 cm dishes of cells, 1-2×10^7^ cells) of the transfected single cells and use the parental cell line as the negative control.

**41.** Prepare five 96well plates for single cell sorting by adding 100 μl growth medium to each well.

**42.** Prepare cells for FACS single cell sorting. Remove growth medium from 10 cm dish of HeLa Kyoto cells. Wash cells with 5-10 ml of PBS per 10 cm dish. Add 1-2 ml Trypsin-EDTA 0.05% and incubate at RT until cells are almost detached from the dish (1-5min). Remove Trypsin-EDTA 0.05% and add 2 ml FACS buffer. Resuspend cells thoroughly and filter cells through a tube with cell strainer. Use HeLa Kyoto cells as negative control for FACS.

**43.** Perform single cell sorting for cells expressing the fluorescent marker (usually mEGFP or mCherry) into 96 well plates containing 100 μl growth medium whereby at least five 96 well plates containing single cells are generated. The fluorescent signal of endogenously tagged proteins are usually relative dim. The efficiency of positively tagged cells is in the range of 0.05-10% positive cells and is typically 0.5%.

**44.** Day 2-12: Let the cell clones grow at 37°C, 5% CO_2_ in complete growth medium for the next 2-12 days. Check after ∼5 days in which wells cells are grown and mark the wells with single cell clones. Single cell clones are growing as a single colony. Discard wells containing multi cell clones which would grow as multi colonies at different areas of the well. Grow the single cell clones so that ∼ 50% of the well are covered by cells.

**45.** Day 10-12: Pick survivors and transfer them onto a fresh 96well plate, so that all clones are now on only one or two 96 well masterplates. To pick the clones, remove the medium and rinse the cells once with 1X PBS. Add 20μl of trypsin into each well and incubate at RT for 1min, remove trypsin, add 100μl of complete growth medium and transfer all cells into a fresh well of a 96 well plate.

**46.** Day 13-14: Split masterplates by trypsinization as described in step 45 and prepare duplicates for DNA preparation.

**47.** Keep clones growing in complete growth medium at 37°C, 5% CO_2_ on 96well plates until you are ready to assess if the clones are expressing the correctly tagged POI. Split the cells by trypsinization at ∼80% confluency.

**48.** *To validate the knock-in cell lines perform the following tests.* Follow option A to perform junction PCR *to test the* fluorescent marker has integrated at the correct locus and test of homozygosity. Follow option B to perform Southern blot analysis to again show homozygosity and to exclude off-target integration of the tag. Follow Option C to perform Sanger sequencing to detect mutations within the insertion site of the tag at the genomic locus. Follow option D to perform Western blot analysis to investigate the expression of randomly inserted tags.

### *Option A.* Validation of fluorescent marker integration at correct locus and test of homozygosity by junction PCR TIMING 1 d

i. *Design of oligos for junction PCR.* Design a forward primer binding at the 5’ end outside of the left homology arm and a reverse primer binding to the fluorescent marker gene (Table 1). This primer set should result in one PCR product of the expected size. To test if all alleles are tagged with the fluorescent marker at the correct locus (i.e. they are homozygously tagged), design a forward primer located 5’ outside of the left homology arm and a reverse primer 3’outside of the right homology arm. Two PCR products using this primer set will indicate heterozygote clones and one PCR product which runs at the expected size of the tagged gene will indicate homozygosity (Supplementary Figure S2).
ii. **CRITICAL STEP** The junction PCR is not a quantitative PCR; consequently, the tagged-gene in heterozygous clones is sometimes not detected as there is a competition between the two PCR products (Figure 4). If problems occur in detecting the tagged gene, it is advisable to repeat the junction PCR with primers used for the T7E1 assay, which result in shorter PCR products, thereby improving amplification of both PCR products. However, using the T7E1 primers has the disadvantage that the primers are localized within the homology arms, which will also amplify randomly integrated donor plasmids. Homozygously tagged genes will always generate one PCR product. Genomic DNA prepared from parental cell lines such as HeLa Kyoto cells are used to detect the untagged gene. Water control is used to detect possible contaminations that can occur during PCR.
iii. *DNA preparation for junction PCR.* Prepare DNA directly from 96well plates by removing the medium from each well.
iv. Add 30 μl DNA lysis buffer.
v. Resuspend cells and transfer the DNA solution into PCR tubes.
vi. Incubate DNA solution at 56°C for 1h followed by 95°C for 10min. PAUSE POINT Store the DNA at 4-8°C until further use.
vii. *Junction PCR.* Amplify the region of interest using the HotStar HighFidelity PCR kit, each reaction was set up as described below:

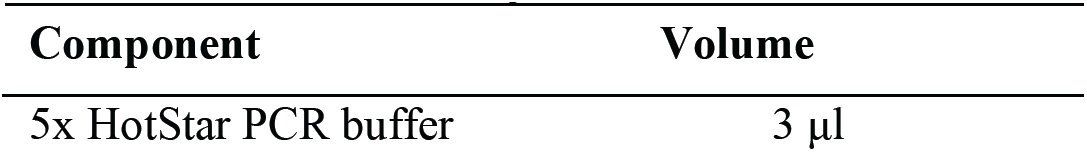
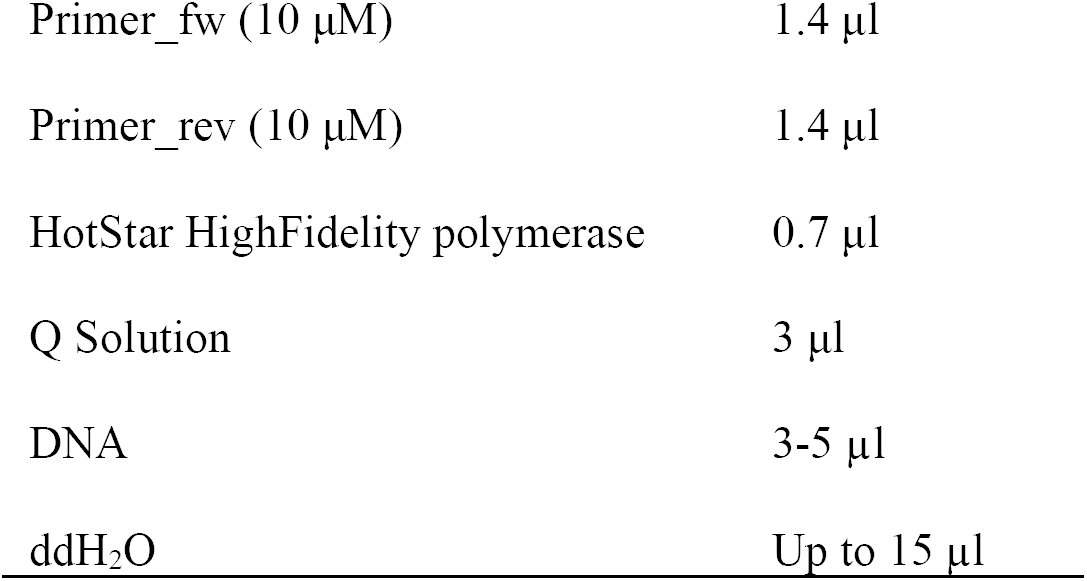 Amplify the region of interest using the following cycling conditions:

viii. *Agarose gel electrophoresis*. Prepare a 0.8% agarose gel using 1X TBE.
ix. Submerge gel into electrophoresis tanks containing 1X TBE.
x. Apply 5 μl of Gene Ruler as marker.
xi. Add 3 μl of 6x Loading dye to the PCR product and apply to a 0.8% agarose gel using 8-channel pipette.
xii. Run samples at 100-150Volt for 60-90min. to gain good separation. Expected result is shown in Figure 4 and Supplementary Figure S2.

Amplify the region of interest using the following cycling conditions:
| Cycles | Denature | Anneal | Extend |
| --- | --- | --- | --- |
| 1 | 95 °C, 5 min |  |  |
| 2-21 | 95 °C, 30s | Start temperature is 5°C higher than T<sub>m</sub> of oligos, - 0.5°C/cycle, 45s | 68°C, 0.5kb/1min. |
| 22-43 | 95 °C, 30s | End temperature of last cycle = 5°C lower than T<sub>m</sub>, 45s | 68°C, 0.5kb/1min. |

### *Option B.* Southern blot analysis TIMING 5 d

**CRITICAL** As a negative control perform the southern blot with the cell clones and genomic DNA prepared from parental cell lines which do not express tagged POI.

**i**. *Southern blot probe design and labelling.* Ensure that the southern blot probe for the GOI is 100-400 bp and the GC% content is less than 40%. Ensure this probe binds outside of the homology arms. In addition, design a probe against the fluorescent marker.

**ii.** Design forward and reverse primers to amplify the probe from genomic DNA using the HotStar HighFidelity PCR kit as described in step 33.

**iii.** Insert the amplified probe into the Zero Blunt TOPO PCR Cloning Kit according to the manufacturer’s protocol.

**iv.** Use the probe-TOPO plasmid to label the southern blot probe using the PCR DIG probe Synthesis kit from Roche according to the supplier’s protocol.

**v.** Run 2 μl of product on a gel next to known quantities of marker to estimate the amount of synthesized probe. The labelled probe migrates less than the unlabeled one PAUSE POINT The labelled probe can be stored at −20°C or can be used directly.

**vi.** Dilute 10ng labelled probe/ml of hybridization buffer (normally 20-35ul of labelled probe in 25 ml hybridization buffer). Hybridization temperature is dependent on the probe and is calculated as follows:

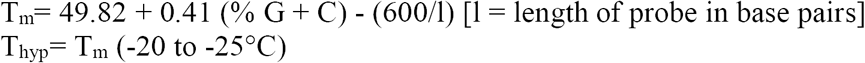

**vii.** *DNA sample preparation.* Prepare genomic DNA using the Epicentre MasterPure DNA purification kit (or a similar genomic DNA purification kit to prepare high concentrated genomic DNA) following manufacturer’s instructions. Two 10 cm cell dishes result in enough DNA for 2-3 Southern blots. Examine 1 μl of genomic DNA on a 0.8-1% agarose gel to detect if the DNA is fragmented and to estimate the amount of DNA.

**CRITICAL STEP** Nano Drop cannot measure genomic DNA precisely.

**viii.** *Digestion of genomic DNA.* Select enzyme(s) such that you will have a detectable difference in size between integrated and non-integrated samples with your probe. Use restriction nucleases that are not sensitive to methylation of CpG dinucleotides and that produce a relatively small fragment. Digest ∼10-20ug of DNA overnight at the appropriate temperature in 40 μl with 1/10v enzyme(s). Load samples on a 0.6-0.8% TBE gel and electrophorese overnight to get a good separation. The genomic DNA of mEGFP-Nup358 was digested with BamHI and SphI resulting in a 3.7kb fragment for the mEGFP-Nup358 and 3kp if Nup358 was untagged (Figure 6).

**ix.** Day 3: *Transfer of DNA onto nylon membrane.*Take a picture of the gel next to a fluorescent ruler and trim off excess gel. Cut the bottom of the right corner for later orientation.

**x.** Wash gel twice for 20min in denaturation solution (∼300-400 ml for a 20 cm x 20 cm gel) on a shaking platform.

**xi.** Rinse the gel once with ddH_2_O and wash 2-3 x 20min with neutralization solution. Check the pH of the gel. Use pH indicator paper and place this onto the edge of the gel to check the pH. The pH should not be higher than 7.5. If the pH is higher, replace the neutralization solution and continue with washes until the pH of the gel is 7.5.

**xii.** Build up southern blot transfer as depicted in Supplementary Figure S3 and as decribed within the following steps xii-xxiii: Fill a tank (20×30 cm) with 2-3L 10 X SSC.

**xiii.** Place a glass plate on top of the tank whereby the glass plate does not cover the whole tank and there is some space in the front and back of the tank.

**xiv.** On top of the glass plate place 2 wicks of 3MM Whatman filter paper which are 2-3 cm wider than the gel and soaked in 10X SSC. Rest the ends of the wicks in 10X SSC.

**xv.** Roll out air bubbles with a plastic pipette.

**xvi.** Soak 1 piece of filter paper the same size as the gel in 10X SSC and place on top of the wicks.

**xvii.** Roll out air bubbles with a plastic pipette.

**xviii.** Slide the gel onto a glass plate and place it onto the wet filter paper.

**xix.** Put saran wrap or parafilm around the edges of the gel on all sides to stop evaporation and short circuits.

**xx.** Cut the Genescreen PlusR nylon membrane to the size of the gel. First pre-wet Genescreen nylon membrane in ddH_2_O and subsequently soak it in 10X SSC. Put the membrane on top of the gel (try not to move it too much!).

**xxi.** Soak 2 pieces of filter paper in 10X SSC and place on top of the membrane, roll with a pipette gently to remove air bubbles.

**xxii.** Place 2 dry pieces of filter paper on top and then stack paper towels on top of the filter paper.

**xxiii.** Place a glass plate on top of the paper towels and two 100 ml bottle half full of water on top as a weight. (the weight should not be too heavy as it will crush the gel). Leave to transfer overnight (>16h).

**xxiv.** The following day, remove the paper towels and paper. Mark the wells of the gel on the membrane using a pencil and turn the damp membrane upright (DNA side is on top.) Keep either some wet filter paper underneath of the membrane or put the membrane onto filter paper soaked in 2X SSC. Crosslink the membrane using a Stratalinker^®^ UV Crosslinker and autocrosslink twice. Afterwards rinse the membrane with ddH_2_O and either continue with hybridization or dry the membrane on filter paper.

PAUSE POINT The dried membrane can be stored at 4°C for at least 1 month.

**xxv.** Day 4: *Hybridization.* Preheat the hybridization oven and pre-warm 50-55 ml DIG Easy Hyb buffer in a water bath at the calculated hybdrisation temperature (see Option B step vi)

**CRITICAL STEP** The hybridization temperature is dependent on the probe (see Option B step vi)

**xxvi.** If the membrane is dry, rinse it briefly with ddH_2_O or 2X SSC. Do not allow the membrane to dry after this point. Place the membrane in the hybridization bottle, by rolling the membrane around a pipette, put the pipette into the bottle and slowly unroll the membrane onto the walls of the bottle. Air bubbles between membrane and glass should be avoided. If the membrane is big and overlaps itself in the hybridization bottle, use a nylon mesh cut 0.5 cm bigger than the membrane, pre-wet it in 2XSSC and roll up with the wet membrane and insert them together into the hybridization bottle.

**xxvii.** Add 25-30 ml pre-warmed DIG Easy Hyb buffer to the hybridization bottle and incubate in hybridization oven whilst rotating for at least 30 minutes.

**xxviii.** Add 15-35 μl probe (final amount should be 125-250ng) into a 1.5 ml eppendorf containing 100 μl ddH_2_O and incubate at 95°C for 5-10min. Subsequently place the denatured probe directly onto ice for 2-3min. Spin probe at RT with 2000x g for 30sec to collect evaporated solution.

**xxix.** Add the denatured probe to freshly pre-warmed 25 ml DIG Easy Hyb buffer.

**xxx.** Exchange the DIG Easy Hyb buffer with the hybridization buffer containing the denatured probe.

**xxxi.** Incubate at the specific hybridization temperature in the oven rotating overnight.

**xxxii.** Day 5: *Detection of probe-target-hybrid.* Wash the membrane twice for 5min in 2X SSC containing 0.1% SDS at RT shaking.

**CRITICAL STEP** All wash steps are done on a shaking platform or with rotation in the hybridization oven.

**xxxiii.** Wash the membrane twice for 15min at 65°C in high stringency wash buffer containing 0.1X SSC and 0.1% SDS (pre-heat wash buffer) with agitation.

**xxxiv.** Wash the membrane once for10min with high stringency wash buffer at RT followed by a 5min wash in DIG1 wash buffer.

**xxxv** Dilute 10X DIG blocking buffer 1:10 with 1X DIG1 buffer.

**xxxvi** Add the 1XDIG blocking solution from step xxxv and block the membrane at RT with rapid agitation for at least 30min.

**xxxvii** Prepare a 1:20000 dilution of anti-digoxigenin-Alkaline Phosphatase Fab fragments in 1X DIG blocking buffer (= anti-DIG solution).

**xxxviii** Remove the 1X DIG blocking solution from the membrane and replace with anti-DIG solution.

**xxxix** Incubate with agitation at RT for 30min.

**CRITICAL STEP** Do not increase this incubation time as background will increase if the incubation time exceeds 30-40min.

**xl.** Wash membrane three times for 20min each with DIG1 wash buffer and twice for 2min with DIG3 buffer with agitation.

**xli.** Drain membrane slightly and place on a plastic sleeve protector (DNA side upwards).

**xlii.** Add 4-8 ml CDPstar chemiluminescent substrate onto the membrane and place cautiously on top the sheet of the sleeve protector avoiding air bubbles.

**xliii.** Incubate at RT 5min.

**xliv.** Remove excess liquid and put the membrane sealed in the sleeve protector into a film cassette.

**xlv.** Place Kodak BioMax MR films directly onto membrane in a dark room. Two films on top of each other can be used to get different exposures at once.

**xlvi.** Expose the films for 1-2h.

**xlvii.** Develop films with the Kodak RPX-OMAT processor or an equal film developer.

**xlviii.** If required, perform a second exposure overnight. PAUSE POINT Remove membrane and store in 2X SSC until further use.

**xlix.** *Stripping and re-probing.* Rinse the membrane 3x in ddH_2_O.

**l.** Incubate the membrane twice for 15min in stripping buffer at 37°C with gentle agitation.

**li.** Wash the membrane twice with 2X SSC at RT for 5min.

**lii.** Use another probe and repeat the hybridization and detection steps xxv-xlviii.

PAUSE POINT Store membrane in 2X SSC at 4°C or

**TROUBLESHOOTING**

### Option C Sanger sequencing TIMING 2 d

**i.** Using genomic DNA (Option B, step vii) prepared with the GenElute Mammalian Genomic DNA Miniprep Kit (Sigma) or a similar genomic DNA preparation kit according to the supplier’s protocol, perform a PCR reaction using the conditions as for the T7E1 assay (step 33) or junction PCR to check for integration of the tag (Table 1) and homozygosity (Option A).

**CRITICAL STEP** Use primers resulting in short fragments, as there are many repetitive sequences and polyA regions within the homology arm that result in poor or disrupted amplification. Furthermore, use a high fidelity proof reading Taq polymerase such as HotStar HighFidelity PCR kit from QIAGEN. Use genomic DNA prepared from parental cells, such as HeLa Kyoto cells, as control for the sequence of the untagged gene.

**ii.** Run the PCR product on an 1-2% agarose gel until good separation of fragments.

**iii.** Gelpurify the PCR products for the tagged and untagged gene using the MinElute Gel Extraction Kit from QIAGEN according to the supplier’s protocol.

**iv.** Perform Sanger sequencing ^41,42^ with primers used for the PCR, to detect possible mutations at the integration site of the tag and additionally, to detect mutations in the untagged alleles (Supplementary Figure S4). Sanger Sequencing services are available from various companies such as GATC and details are found for example on the GATC home page (https://www.gatc-biotech.com/en/index.html).

**v.** If only heterozygously tagged genes were generated, sequence the untagged alleles to find wildtype alleles to perform a second round of transfection with the aim to generate homozygously tagged genes.

**TROUBLESHOOTING**

### Option D. Western blot analysis TIMING 3 d

**i.** *Preparation of protein lysates.* Harvest cells from a 6well cell culture dish by trypsinization and transfer the cell suspension into 2 ml Eppendorf tubes. Prepare also protein lysates derived from parental cells to compare expression levels of untagged versus tagged protein.

**ii.** Pellet cells at 300 x g at RT for 1min.

**iii.** Wash cell pellet with 1X PBS. Spin the cell suspension at 300 x g at RT for 1min. and discard the supernatant.

**iv.** Resuspend the cell pellet well with an equal volume of protein lysis buffer.

**v.** Incubate cell lysate on ice for 30min and mix every few minutes by vortexing or pipetting.

**vi.** Centrifuge the cell suspension at maximum speed (16000 x g) at 4°C for 10min.

**vii.** Determine the total protein concentration of the lysate using the Bio-Rad Protein Assay following the supplier’s protocol.

**viii.** *SDS PAGE.Mix* 40 μg of total protein with 4X NuPAGE^®^ LDS Sample Buffer supplemented with 100 μM DTT.

**ix.** Incubate samples at 65°C for 5-10min.

**x.** Apply sample to 4-12% NuPAGE^®^Bis-Tris Gels in a EI9001-XCELL II Mini Cell (NOVEX) gel chamber and run at 125-150V for approximately 90min. in NuPAGE MOPS SDS Running Buffer (1X, diluted in ddH_2_O). Use PageRuler Plus Prestained Protein Ladder 10-250 kDa as a marker.

**xi.** *Protein blotting.* The protein gel was transferred onto a PVDF membrane using the Mini Trans-Blot^®^ Electrophoretic Transfer Cell.

**xii.** Prepare the transfer buffer.

**xiii.** Cut the PVDF membrane and 3MM Whatmann filter paper to the size of the gel.

**xiv.** Wet the PVDF membrane with methanol, rinse with ddH_2_O and soak in transfer buffer for more than 2min.

**xv.** Assemble the gel sandwich for the transfer as follows: Place the transfer cassette with the black side down, put pre-wetted fiber pads and 2 pre-wetted filter papers on the cassette. Place PVDF membrane on top of filter papers and put gel on top of the membrane. Place two pre-wetted filter papers on top of the gel and roll with a pipette gently to remove air bubbles. Place pre-wetted fiber pad on top and close the transfer cassette.

**xvi.** Place the transfer cassette in the transfer cell such that the membrane site (black side) is placed towards the Anode (+).

**xvii.** Run the transfer in 1 liter of transfer buffer at 4°C with 400mA for 2h or at 60mA overnight.

**xviii.** *Immunodetection*. Disassemble the transfer cassette

**xix.** Block the blots with 5% milk (w/v) in PBS+ 0.1% TWEEN^®^ 20 (= PBS-T) at RT for 1h or at 4°C overnight shaking.

**CRITICAL STEP** perform all incubation and washing steps on a rocker.

**xx.** Dilute the primary antibodies in 5% milk/PBS-T according to Table 2

**Table 2:** Primary antibodies

| **Name** | **Dilution** | **Supplier/ Reference** |
| --- | --- | --- |
| **Anti-GFP (mouse)** | 1:1000 | Roche |
| **RFP antibody [3F5] (mouse)** | 1:500 | ChromoTek |
| **Anti-GAPDH (GC5) sc-32233** | 1:2000 | Santa Cruz Biotechnology |
| **Cas9-HRP antibody ab202580** | 1:2000 | Abcam |

**Table 3:**
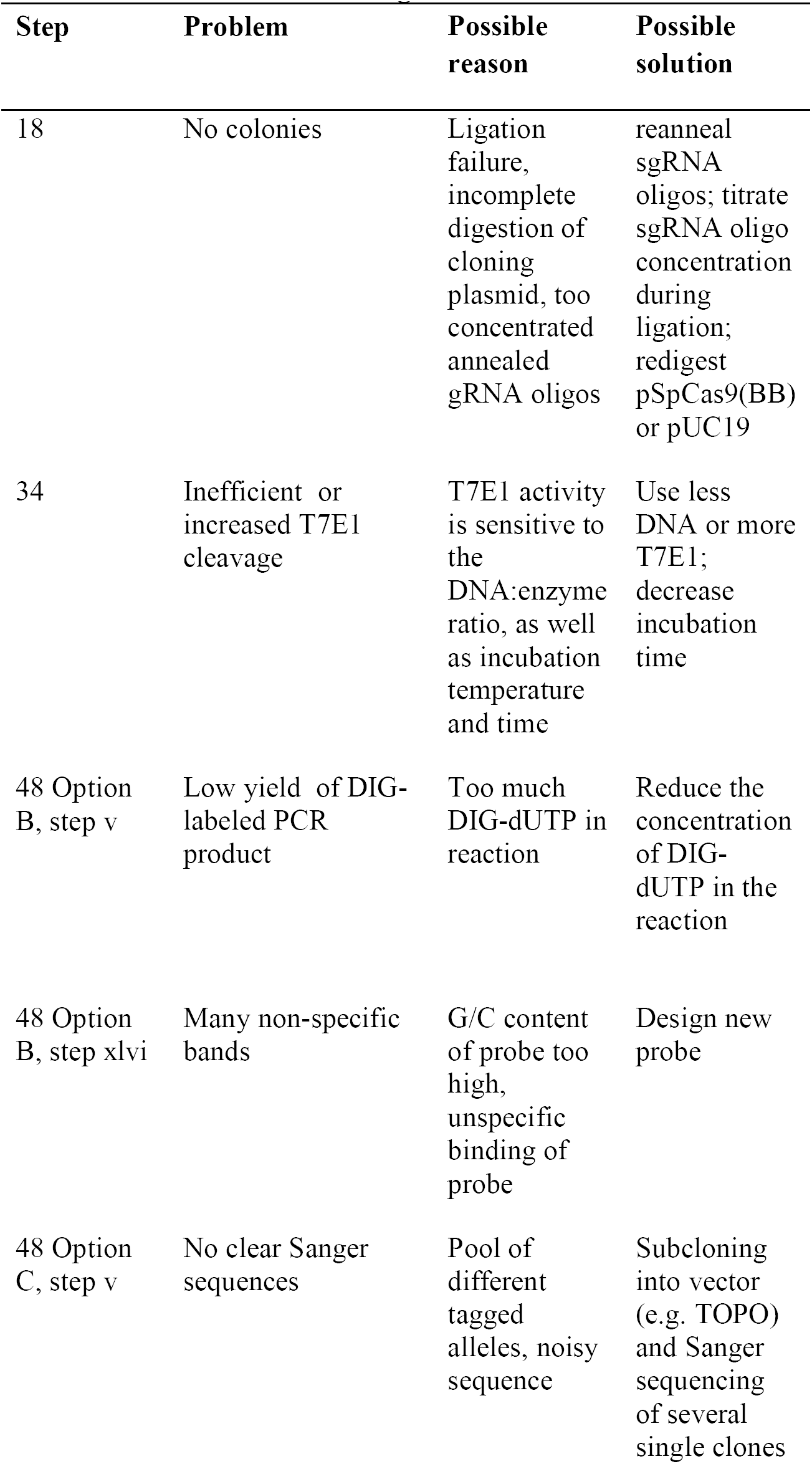
Troubleshooting

**xxi.** Incubate the blot in diluted primary antibody at 4°C overnight.

**xxii.** Wash the blots three times for15 min each with PBS-T at RT.

**xxiii.** Incubate with secondary antibodies diluted 1:5000 in 5% milk/PBS-T at RT for 1h.

**xxiv.** Wash four times for 20min each with PBS-T at RT.

**xxv.** Incubate with Western LIGHTNING ™ *plus*-or with Amersham ™ ECL ™ Prime Western Blotting Detection Reagents according to the manufacturer’s instructions.

**xxvi.** Place Kodak BioMax MR or Amersham Hyperfilm MP films on top of the blots (protein site up) and develop them in a Kodak RPX-OMAT processor (http://www.frankshospitalworkshop.com/equipment/documents/x-ray/user_manuals/x-ray_film_processor/Kodak_X-Omat_M6B_-_User_manual.pdf).

**xxvii.***Stripping of blots* Rinse blots with ddH_2_O and wash for another 5 min in ddH_2_O. Incubate blots for 5-10min. in 0.2 N NaOH.

**xxviii.**Rinse the blots with ddH_2_O and wash them for 5min in ddH_2_O.

**xxix.** Then perform the immunodetection protocol with another primary antibody.

**CRITICAL STEP** Use antibodies against the POI if available to detect homozygous expression of cell clones. Immunoprecipitations of the most promising clones are performed to test for the integration of the tagged protein in the expected complex.

### Microscopy of promising tagged cell clones TIMING 2 d

**49.** To detect correct localization of the tagged protein, follow option A to perform live cell imaging or option B to perform immuno-staining with fixed cells. To perform cell cycle analysis of the tagged cell clones see Box 1.

**(A) Live cell imaging**

**CRITICAL** Live cell imaging has the advantage to follow a cell throughout the cell cycle and investigate if the tagged protein is performing as expected during cell division. Prior to live imaging set up the correct cell density and growth conditions on imaging plates. If cells do not grow on glass bottom plates, use plastic imaging plates.

**i.** Seed all cell clones, which have been positively tested for the integration of the fluorescent marker at the correct locus, onto a glass bottom 96well plate for imaging. The cell density should be 50-70% during imaging, this is usually 5000-7000 HeLa Kyoto cells per 96well.

**ii.** Grow cells in normal growth medium at 37°C, 5% CO_2_ overnight.

**iii.** Change medium to CO_2_-independent medium without Phenol red and seal the edges of the plate with silicon.

**iv.** Use a confocal microscope such as the Zeiss 780 laser scanning confocal microscope for live cell imaging. Maintain cells growing at 37°C in an incubation chamber within the microscope. Take images with a 63× PlanApochromat numerical aperture (NA) 1.4 oil objective during the cell cycle to observe if the fluorescently tagged protein behaves as expected.

**(B) Imaging of fixed cells**

**CRITICAL STEP** Imaging of fixed cells has the advantage of allowing counter staining with antibodies if necessary. The correct localization of the tagged POI can be confirmed by the use of antibodies against the POI.

**i.** Plate cells as described in step 49 option A on glass bottom plates or any other imaging plates which can be used on confocal microscopes.

**ii.** Grow the cells in normal growth medium at 37°C, 5% CO_2_ overnight.

**iii.** Fix the cells with 4%PFA in 1X PBS at RT for 15min.

**iv.** Wash with 1X PBS twice.

**v.** If necessary counter stain nuclei with 0.1 μg/ml HOECHST in PBS.

**vi.** Keep cells in 1X PBS for imaging.

**PAUSE POINT** Fixed samples can be stored at 4-8°C for 1-2 weeks.

**vii.** Perform imaging as described in step 49 option A. Try to image them within 1-2 days after fixation. However imaging up to 2 weeks can be performed, but can result in decreased signals.

### Box 1 Cell cycle analysis TIMING 4 d

**CRITICAL** Establish the best growth conditions and seeding density for the cell line to be analyzed. Cells should grow evenly, with a confluency of ∼50% at the beginning of imaging. Adjust growth conditions for each cell line. If necessary use CO_2_ and normal growth medium w/o phenol red for imaging. Any wide-field microscope with incubator can be used as long as it is equipped for autofocus and automated imaging (e.g. ImageXpress Micro XLS Widefield High-Content Analysis System from Molecular Devices or ScanR from Olympus). The conditions below are best for HeLa Kyoto cells. The cell cycle of the endogenously tagged cell lines are compared to the cell cycle timing of the parental cells such as HeLa Kyoto cells (Figure 8 and 9).

**1.** *Seeding of cells.* Plate 7000 HeLa Kyoto cells per well on a 96Well SCREENSTAR Microplate. Grow cells at 37°C and 5% CO_2_ overnight.

**2.** Day 2: Stain the nucleus with 50nM SiR-DNA in CO_2_-independent imaging medium (w/o Phenolred) and seal the edges of the plate with silicon. Incubate at 37°C (no CO_2_) for 3h-4h.

**3.** Day 2-3: *Imaging.* Take images in the Cy5 Channel every 5 minutes for 24h using ImageXpress Micro XLS Widefield High-Content Analysis System (Molecular Devices). Perform an autofocus for all imaged wells and set up the imaging conditions so that the signal to noise ratio is 3:1.

**4.** Day 4: *Cell cycle analysis.* Use the CellCognition software^47 (^CeCogAnalyzer 1.5.2) to automatically track single cells through the cell cycle and classify them into interphase and mitotic stages.

**CRITICAL STEP** The code and a general documentation for the cell cycle analysis are available on GitHub via https://git.embl.de/grp-ellenberg/Cecog_HMM_PostProcessing and the related wiki https://git.embl.de/grp-ellenberg/Cecog_HMM_PostProcessing/wikis/home.

The settings and documentation for CellCognition can be found on GitHub under the following link: https://git.embl.de/grp-ellenberg/Cecog_HMM_PostProcessing/wikis/cellcognition

The CellCognition output is further processed with the custom-written MatLab code *cellfitness.m* to determine the mitotic timing including statistical analysis. The documentation for Cellfitness can be found on GitHub under the following link: https://git.embl.de/grp-ellenberg/Cecog_HMM_PostProcessing/wikis/cellfitness.

## TIMING

Steps 1|-3|: gRNA design: 1d

Steps 4|-5|: Donor design and synthesis: 1d design, 13-20d waiting time

Steps 6|-18|: Cloning gRNA into vector: 4d consisting of 1-2h per day hands on, 2x overnight incubations and 1d waiting of sequencing results

Steps 19|-37|: Functionality test of gRNAs: 5d in total consisting of ∼1h hands per day and 2-3d incubation

Steps 37|-40|: Delivery of paired CRISPR/Cas9: 7-10d in total consisting of 0.5h hands on at day 1+2 and day 2-10 is waiting time.

Steps 41|-47|: Selection of single cell clones via FACS: 10-12d in total consisting of 1 d sorting and 10-11 days waiting time

Step 48 Option A|: Junction PCR: 1 d consisting of 1-2h hands on and 6h waiting time

Step 48 Option B |: Southern blot: 5 d in total consisting of 1-2h hands on each day and 3x overnight waiting time

Step 48 Option C |: Sequencing: 2d consisting of 1-2h hands on and 1-2 waiting days

Step 48 Option D |: Western blot: 3d in total consisting of 1-2h hands on each day and 2x overnight incubation

Steps 49 Option A and B|: Microscopy: 2d consisting of 0.5-1h hands on per day and 1x overnight incubation

Box 1: Cell cycle analysis: 4d in total consisting of 1h hands on at day 1+2, 24h automated imaging and 1d data analysis

## ANTICIPATED RESULTS

70% of the targeted genes were tagged homozygously with FPs and resulted in physiological levels and phenotypically functional expression of the fusion proteins as demonstrated representatively with HeLa Kyoto mEGFP-NUP358 cells in this protocol. In general, homozygous tagging is not feasible for some proteins when structure and size of the FP tag perturb the function of the fusion protein. Therefore, some genes can only be tagged heterozygously. Furthermore, it should also be considered how many molecules of the POI are expressed because there are limits for some applications when expressing low amounts of tagged proteins.

During the validation pipeline there will be some cell clones that do not behave like parental cell lines and these clones are eliminated for further studies.

However, the big advantage of endogenously tagging is that there will not be any overexpression artefacts^1^ keeping physiological expression levels and functionality of the tagged POI. The anticipated results of each step of the validation pipeline are outlined below: The functionality test of gRNAs of the negative controls will result in one band of the same size as the PCR product after the T7E1 digestion whereas the paired nickase sample digested with T7E1 should have several cleavage bands as predicted. The gRNA pair with the highest cleavage efficiency will be used for introducing the fluorescent marker gene at the locus of interest.

The expression levels of the endogenously tagged protein are low during the selection of single cell clones expressing the fluorescent tag via FACS. Additionally, the efficiency of positively tagged cells 7-10 days after plasmid transfection ranges between 0.05-10%. Therefore, it is essential to use HeLa Kyoto wt cells as negative control to distinguish between non-specific and specific FP expression levels. Figure 3 displays a typical FACS result.

The junction PCR is performed in general with two primer sets. To test if the fluorescent marker is integrated into the expected locus, a forward primer binding at the 5’ end outside of the left homology arm and a reverse primer binding to the fluorescent marker gene (Table 1). This primer set should result in one PCR product of the expected size. To test if all alleles are tagged with the fluorescent marker at the correct locus (i.e. to test homozygosity), use a forward primer located 5’ outside of the left homology arm and a reverse primer 3’outside of the right homology arm. Two PCR products using this primer set will indicate heterozygous clones and one PCR product which runs at the expected size of the tagged gene will indicate homozygosity (Supplementary Figure S2). A typical result of the junction PCR is shown in Figure 4. This is not a quantitative PCR, therefore the tagged gene in heterozygous clones might not be detected due to competition between the two PCR products (Figure 4B). Therefore, it is advisable to repeat the junction PCR with primers within the homology arms resulting in shorter PCR products improving amplification of both PCR products demonstrated in Figure 4C.

Southern blot analysis is used to detect homozygosity, additional integration of the tag and/or rearrangements within the endogenous locus of interest. In Figure 6 a typical result of a Southern blot analysis is depicted. Homozygous clones result in one band detected with both probes because only the tagged gene is present (Figure 6, clone #97). In heterozygous clones one band using the probe against the tag is detected and two bands with the probe against the endogenous locus because the tagged and untagged gene is present (Figure 6, e.g. clone #118). Moreover, extra unexpected DNA fragments, patterns or sizes can be detected due to extra integration of the donor plasmid mutations of the region of interest (Figure 6, e.g. clone #1). Clones showing unexpected DNA fragments or unexpected pattern or sizes are not used for further experiments.

Sanger sequencing of the cell clones is important to detect possible mutations which were not detected via Southern blot analysis. A sequencing example followed by alignment analysis is shown in Figure 5 and in detail in Supplementary Figure S4. Homozygous clones such as #97 in Figure 5 contain only the tagged mEGFP-NUP358 whereby the mEGFP is inserted correctly into the target site. Mutations can be detected as shown in Figure 5 clone #118 which contains a deletion of 63nt causing in turn a deletion of 6 amino acids in the first exon of the untagged NUP358.

A specific antibody against the POI is very useful to distinguish homo- and heterozygous clones which can be detected via Western blotting. Homozygously tagged POI should solely express the tagged POI resulting in one specific band corresponding to the expected molecular weight as shown in Mahen *et al*.^1^. An antibody against the FP, such as anti-GFP, detects specific expression of the tagged proteins of interest as a specific band at the expected size (Figure 7). If there are additional bands detectable, which are elevated in comparison to background levels in the parental cell lines, either mutations have occurred or free FP is expressed (Figure 7, clones #122 and #26). Clones with unexpected bands should not be used. Live cell imaging as depicted in Figure 10 is used to detect correct localization of the tagged protein. The correct localization of the tagged POI can be confirmed by the use of antibodies against the POI.

**Figure 10:**
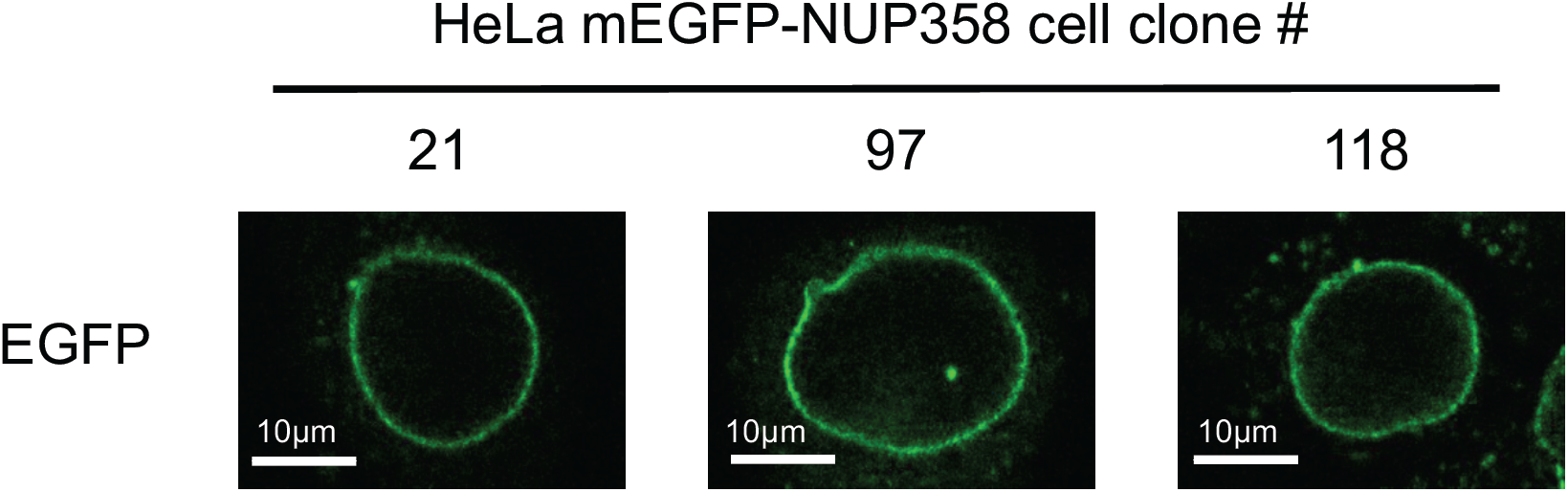
Live cell imaging of HeLa Kyoto mEGFP-NUP358 cell clones. Various HeLa cell clones expressing HeLa Kyoto mEGFP-NUP358. #97 represent a homozygously tagged mEGFP-NUP358 clone whereas #21 and #118 are heterozygously tagged ones.

The automated workflow of the cell cycle analysis including representative results are shown in Figure 8 and 9. The duration of mitosis is compared to wt cells and clones with extended (Figure 8D, mEGFP-CDC20 #140) or shortened mitotic time (Figure 8D, mEGFP-Bub3 #74) are discarded.

To detect correct localization of the tagged protein, live cell imaging or immuno-staining in fixed cells can be performed. Live cell imaging has the advantage of fully preserving the FP fluorescence being able to follow a cell throughout the cell cycle and investigate if the tagged protein localizes as expected at different cell cycle stages.

NUP358 is a nuclear pore complex protein which localizes to the nuclear envelope (NE) during interphase and to annulate lamella which are common in tumor cell lines. HeLa Kyoto mEGFP-NUP358 cell clones were analyzed using live cell imaging and only clones which showed correct localization without additional non-specific GFP localization were selected for further experiments (Figure 10). Clone #97 has additionally to the NE localization also a spot within the nucleus visible, which derived from mEGFP-NUP358 expressed in annulate lamella, that in turn also demonstrates correct localization of mEGFP-NUP358.

## ACKNOWLEDGMENTS

We thank the mechanical and the electronics workshop of EMBL for custom hardware, the Advanced Light Microscopy Facility of EMBL for microscopy support and the Flow Cytometry Core Facility of EMBL for cell sorting. We gratefully acknowledge G. Reid for critically reading the manuscript. This work was supported by grants to J.E. from the European Commission EU-FP7-Systems Microscopy NoE (Grant Agreement 258068), EU-FP7-MitoSys (Grant Agreement 241548), iNEXT (Grant Agreement 653706), as well as by the European Molecular Biology Laboratory (B.K., B.N., M.K., Y.C., N.W., J.E.). N.W. and Y.C. were supported by the EMBL International PhD Programme (EIPP).

## AUTHOR INFORMATION

### Contributions

B.K. designed and performed the experiments. B.K. with the help of B.N., Y.C. and N.W. developed the protocol. B.N. and M.K. tested the protocol. Y.C. and N.W. created the automation of the cell cycle analysis. B.K. with the help of B.N., Y.C., N.W. and J.E. wrote the protocol. All authors contributed to the interpretation of the data and read and approved the final manuscript.

## Supplementary Figures

**Supplementary Figure S1:**
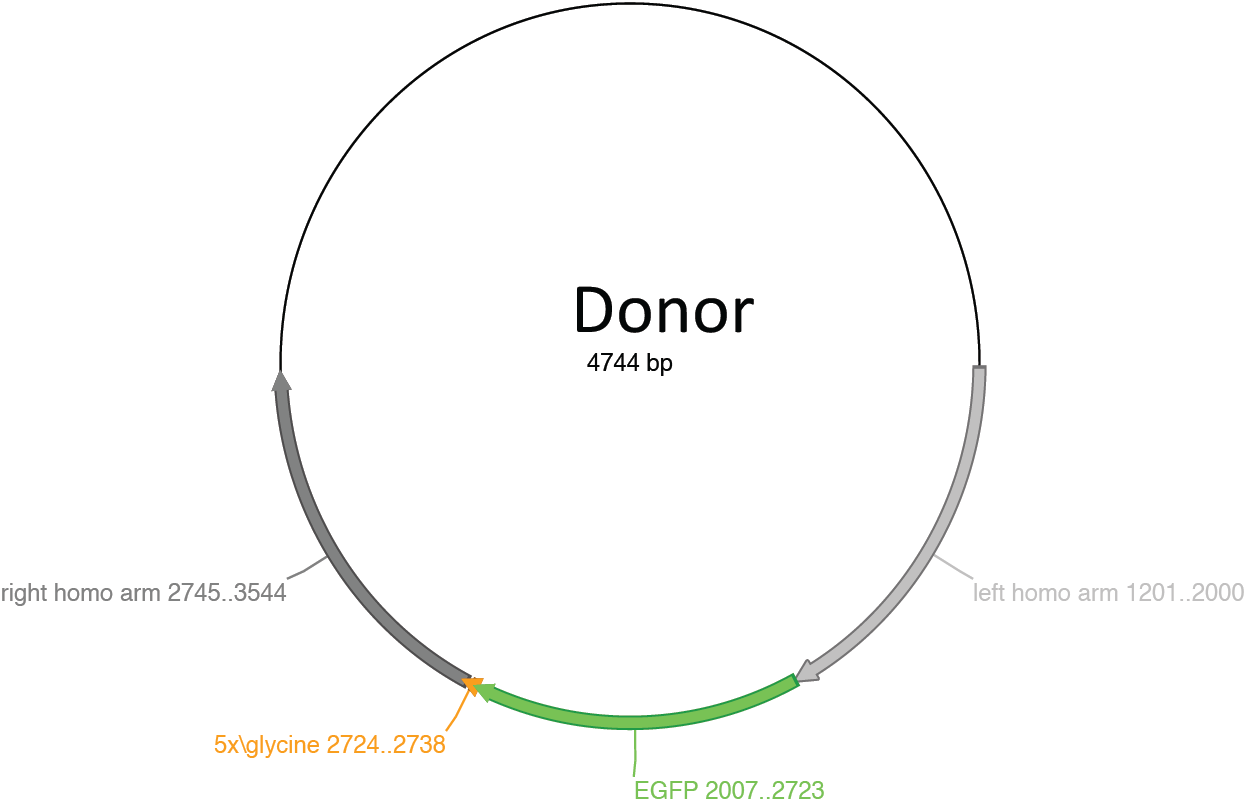
Donor plasmid. The donor plasmid consists of the fluorescent marker gene (mEGFP in this case) which is flanked by 500-800 bp homology arms of the GOI. A linker is placed between the tag and the gene to maintain functionality of the tagged protein. As a backbone vector pUC-based plasmids are used.

**Supplementary Figure S2:**
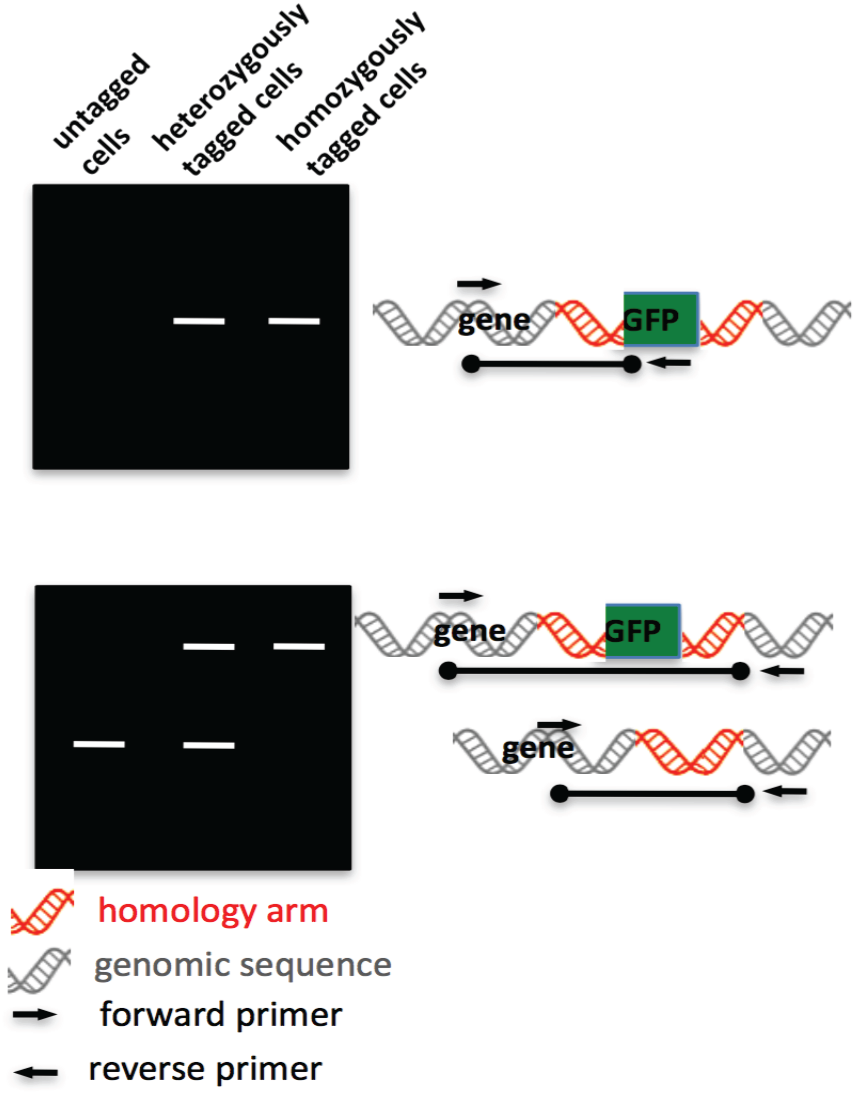
Scheme of expected junction PCR result. A forward primer binding at the 5’ end outside of the left homology arm and a reverse primer binding to the fluorescent marker gene will result in one PCR product of the expected size. To test if all alleles are tagged with the fluorescent marker at the correct locus, a forward primer located 5’ outside of the left homology arm and a reverse primer 3’outside of the right homology arm were used. Two PCR products using this primer set will indicate heterozygote clones and one PCR product which runs at the expected size of the tagged gene will indicate homozygosity.

**Supplementary Figure S3:**
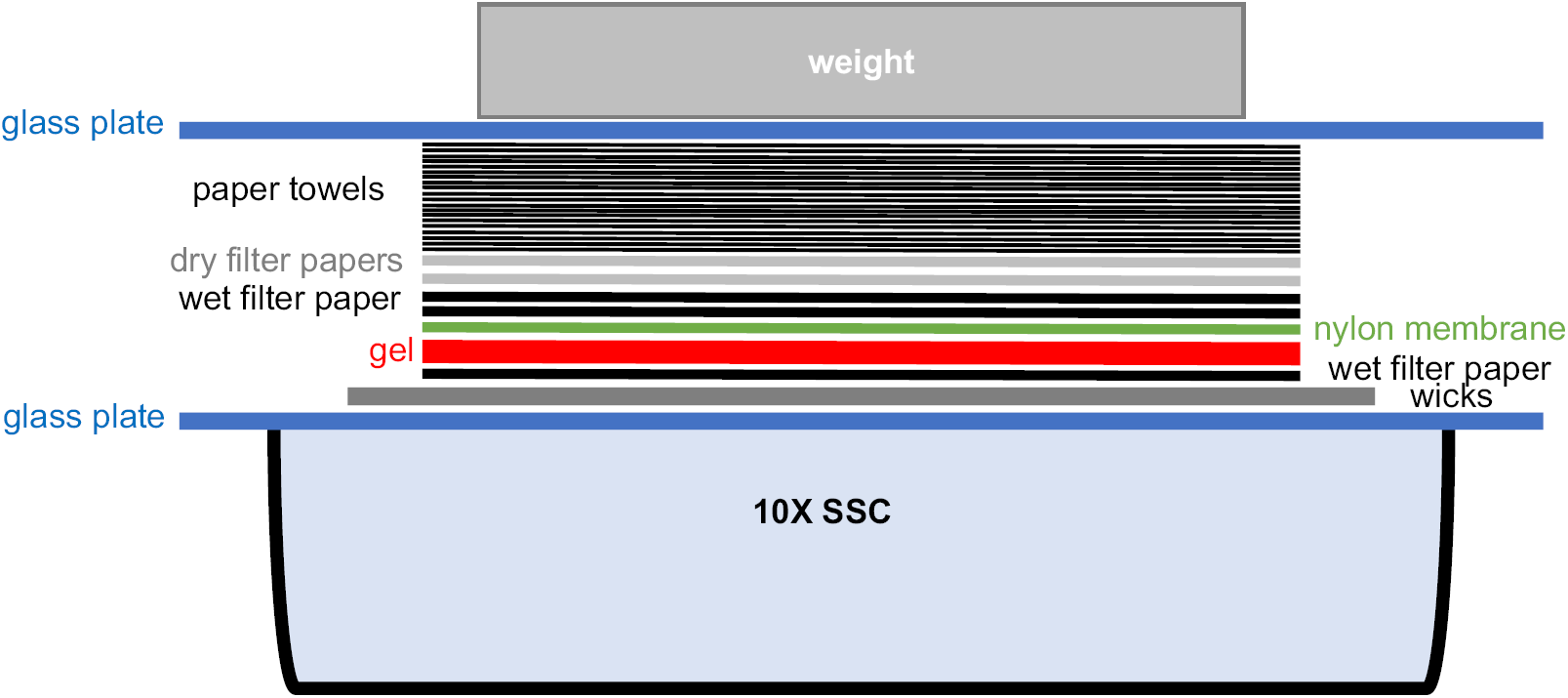
Southern blot transfer of DNA. This scheme depicts how to build up the southern blot transfer as described in step 48| Option B, step xii-xxiii). Two long filter papers are dipped into a tank filled with 10X SSC and used as wicks. Gel, nylon membrane and filter papers were sandwiched on a glass plate as depicted to transfer the DNA from the gel onto the membrane.

**Supplementary Figure S4:**
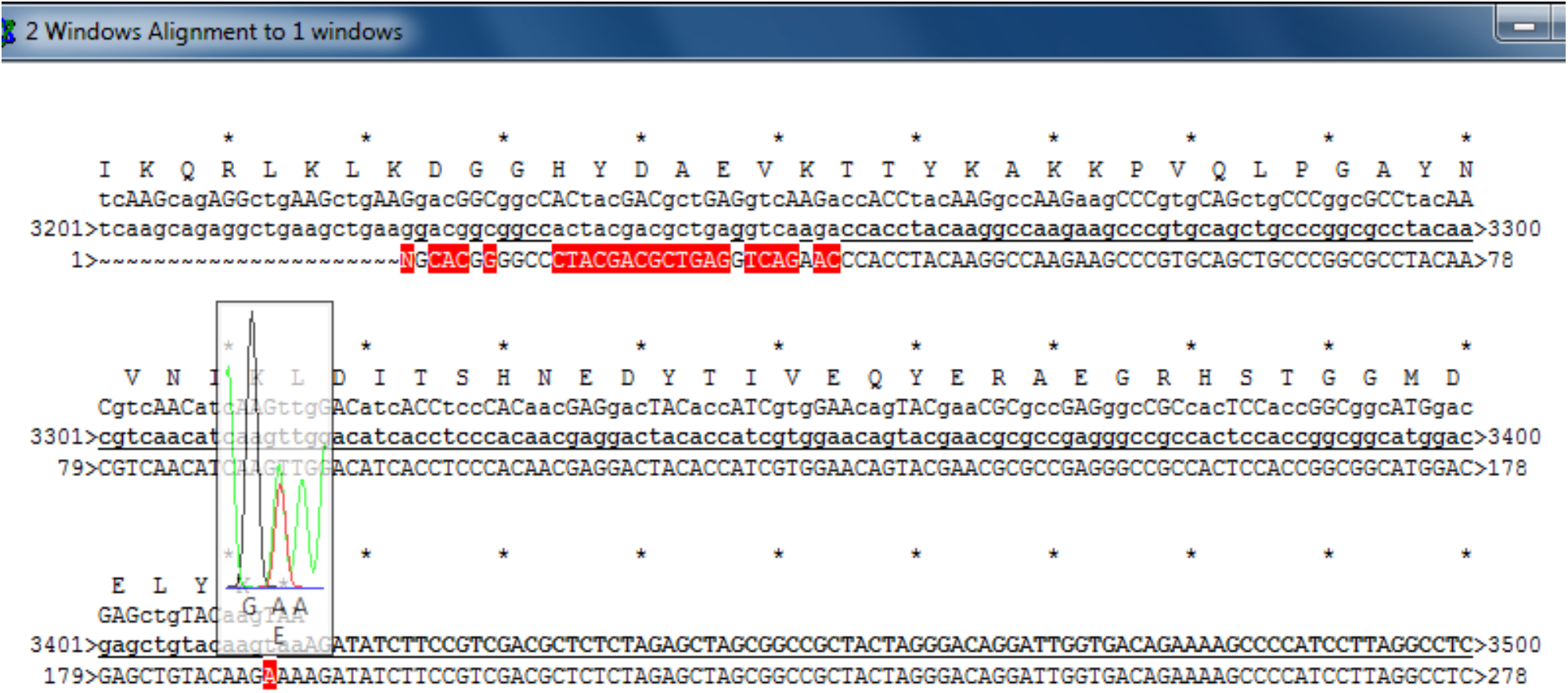
Example of sequences containing point mutation. Sanger sequencing was performed with a PCR fragment (lower sequence line #1-278) which was aligned to the expected sequence of the GOI (middle sequence line labeled with #3201-3500). The overlaying green and red lines indicate a point mutation at position 3411 of the expected sequence, i.e. one allele has the nucleotide T at this position (green line) whereas another allele has the nucleotide A (red line) at the same position. This indicates a point mutation (red A) within one of the alleles.

